# The activator domain of bacterial collagenases drives collagen recognition, unwinding and processing

**DOI:** 10.1101/2023.07.18.549520

**Authors:** Jamil Serwanja, Alexander C. Wieland, Astrid Haubenhofer, Hans Brandstetter, Esther Schönauer

## Abstract

Collagens form the resilient backbone of the extracellular matrix in mammals. Only few proteases are able to digest triple-helical collagen. Clostridial collagenases can efficiently process collagen. However, little is known about the mechanism of bacterial collagenolysis of either soluble collagen or the multi-hierarchically assembled, insoluble collagen fibers. Here we present a functional analysis of the distinct roles of the individual domains of collagenase G (ColG) from *Hathewaya histolytica.* A broad array of biochemical, biophysical, and enzymatic data consistently revealed unexpected synergistic and antagonistic interactions between the activator, peptidase and collagen-binding domains. We found the non-catalytic activator domain to act as a master regulator, coordinating the complex interactions to specifically recognize and process the diverse physiological substrates. The results presented here enable multiple applications such as the engineering of collagenase variants with selectivity for defined substrate states.

## Introduction

Collagens are the most abundant and long-lived proteins in mammals, constituting up to 90% of the extracellular matrix. They are predominantly found as fibrillar type I collagen (∼90%)^1–3^. Its supramolecular fibrils shape the extracellular matrix and are essential for tissue integrity ^4,5^. Soluble monomeric type I collagen (tropocollagen) is a tightly packed right-handed triple helix composed of three α-chains. The triple helix is stabilized by repetitive Gly-X-Y triplets, in which X and Y are frequently occupied by proline and hydroxyproline, respectively ^6^. Triple-helical collagen is highly resistant to proteolysis due to (i) its tight helical packing, caching peptide bonds in the helix interior, and (ii) its high content of (hydroxy)prolines, which only few proteases can accommodate in their active sites ^7,8^. Moreover, fibrillar collagen adopts a complex multi-hierarchical structure. Soluble tropocollagens (∼300 nm length, ∼1.5 nm diameter) self-assemble into larger insoluble complexes, from microfibrils (diameter of > 5 nm) via fibrils (30 – 500 nm diameter) to 0.5-2 µm thick fibers. These fibers are further stabilized by pyridinoline crosslinks, and glycosaminoglycan interactions ^5,9–11^.

A true collagenase must therefore be able to (i) bind to fibrillar collagen, (ii) extract a single tropocollagen from the complex, (iii) unwind the triple helix to excavate the scissile bonds, (iv) accommodate the imino acid-rich α-chain in its active-site cleft, and (v) cleave pre- or post-(hydroxy)proline peptide bonds. Only a small number of mammalian enzymes are capable of this task under physiological conditions and typically, they do so with narrow substrate specificities (*e.g.* MMP-1/2/8/13/14/18, neutrophil elastase) ^12,13^. However, bacteria have also evolved collagenases, most notably *Clostridium* spp., *Bacillus* spp. (M9B subfamily) and *Vibrio* spp. (M9A subfamily) ^14^. In contrast to most mammalian collagenases, these bacterial enzymes are capable of breaking triple-helical collagen down into small peptides ^15–17^. This enables saprophytic species to feed on collagen and provides pathogenic strains with a tool for host infection ^18–21^. However, the molecular mechanisms that enable bacterial collagenases to fully degrade collagen are not completely understood.

Bacterial collagenases are multi-domain zinc-metalloproteases ^15,22–27^. They harbor an N-terminal collagenase unit (CU) of ∼80 kDa, consisting of an activator domain (M9N domain) and a peptidase domain (peptidase M9 domain) connected via a 9 aa-long linker ^23,28,29^. At the C-terminus, a varying composition of accessory domains, *i.e.*, polycystic-kidney disease-like domain(s) (PKD), and collagen-binding domains (CBD) (each ∼10 kDa), is found ^23,24,26,30^ (**Fig. 1a**). The best studied bacterial collagenases are ColG and ColH from *Hathewaya histolytica* (formerly *Clostridium histolyticum*) ^15,21,23,24,31,32^. The high-resolution structure of the CU of ColG revealed an ‘open’ saddle-shaped CU architecture (**Fig. 1b**) and we identified the CU as the minimal collagenolytic entity *in vitro* ^23^. Based on these findings, we proposed a conformational two-state model of bacterial collagenolysis, *aka* chew-and-digest model, in which the processing of triple-helical collagen is driven by the opening and closing of the CU. In this model, tropocollagen is recognized by the PD in the (crystallized) open CU state, thereby triggering CU closing. In the (proposed) compact state, the AD and PD are able to interact with triple-helical collagen, thereby facilitating the unwinding of bound collagen and enabling the cleavage of entrapped α-chains in the (semi-) closed CU conformation. On a more speculative level, the chew-and-digest model was extended to the processing of microfibrils. Hereby, an initially open CU binds a microfibril which triggers a CU contraction, by which a single tropocollagen is exposed from the fibrillar ensemble, unwound and then cleaved by the engulfing CU ^23^.

**Fig. 1:**
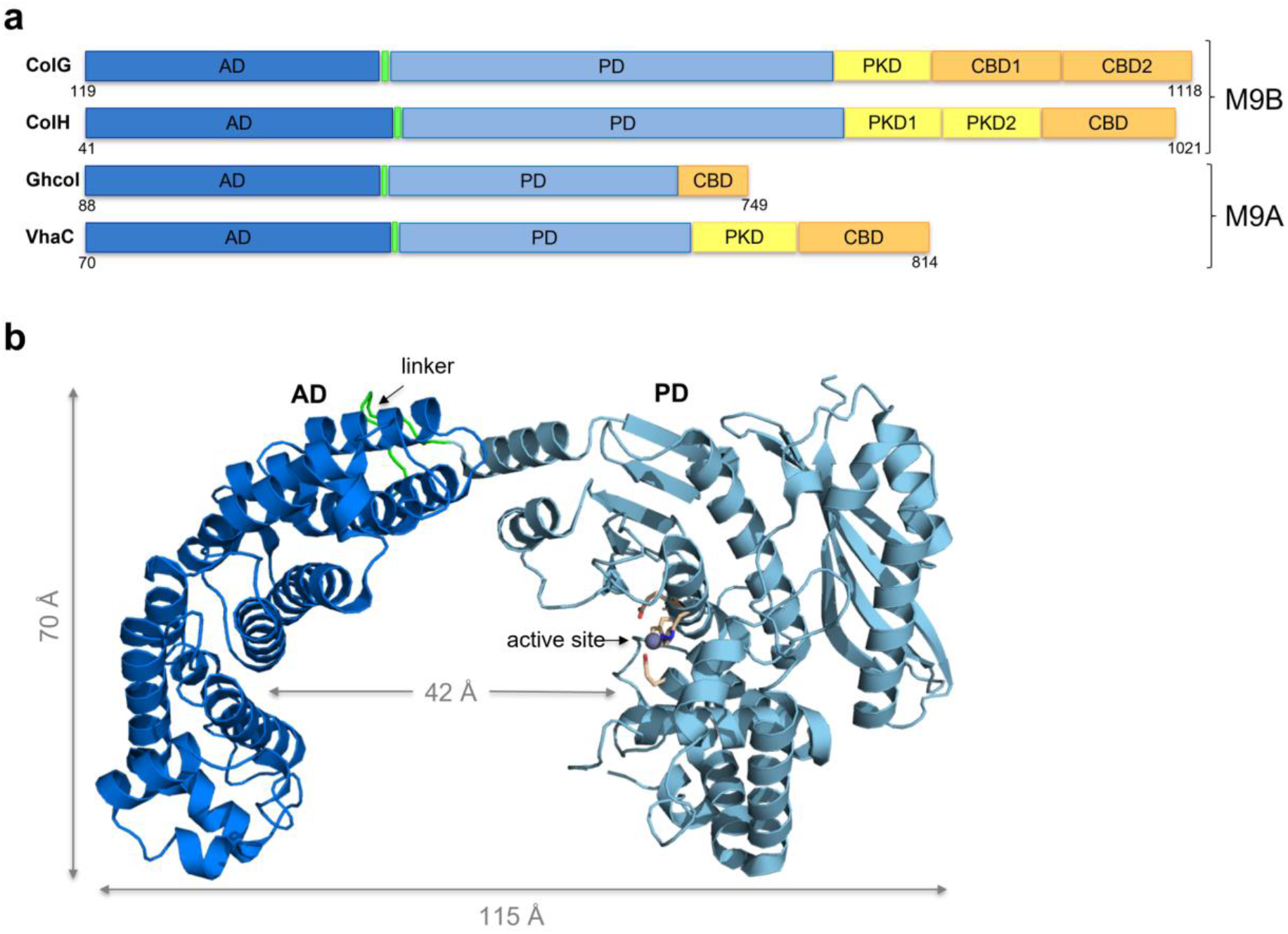
Domain composition and crystal structure of ColG-CU. **a**, Schematic domain organization of ColG and ColH from *H. histolytica*, Ghcol from *Grimontia hollisae*, and VhaC from *Vibrio harveyi VHJR7* ^29,34,58^. Activator domain (AD) (dark blue), linker (light green), peptidase domain (PD) (light blue), polycystic disease-like domain (PKD) (yellow), and collagen-binding domain (CBD) (orange). **b**, Ribbon representation of the collagenase unit of ColG (adapted from Eckhard *et al*. ^23^). The catalytic zinc ion (grey) and the catalytic residues (light orange) are shown in ball-and-stick representation. The linker is highlighted (green). Molecular figures were created with PyMOL^59^.

Support for the existence of the closed CU conformation was provided by a SAXS model of ColH^33^, and crystallographic studies identified similar open saddle-shaped CU architectures of the *Vibrio* collagenases Ghcol of *Grimontia hollisae* ^28^ and VhaC of *Vibrio harveyi VHJR7* ^29^. Even though these findings corroborate two pillars of the chew-and-digest model, namely the existence of an open and (semi-) closed CU conformation, other results argue for a refinement. *Wang et al.* found that collagen was bound by the VhaC-AD, while VhaC-PD showed only minor or no binding affinity. The authors, therefore, suggest that the initial recognition of tropocollagen is mediated by the AD in bacterial collagenases ^29^. However, the essential questions of how bacterial collagenases manage to unfold the collagen triple helix – a feature, which sets them apart from other peptidases - and how they are able to process higher-order assemblies of collagen, are still not answered.

Here, we investigate the role of individual domains of bacterial collagenases in the recognition of the different physiological forms of fibrillar collagen. We identify and characterize the triple-helicase entity in bacterial collagenases, and we demonstrate the significance of AD-PD interplay in the processing of soluble and insoluble collagen using conformational trapping.

## Results

### Bacterial collagenases cleave collagen only after being unwound into 𝛼-chains

Fibrillar collagen adopts a multi-hierarchical structure in the body. Yet, the active-site dimensions of *Clostridium* ^23,34^, *Vibrio* ^28,29^ and mammalian collagenases like MMP-1 ^35,36^ are all around ∼10 Å wide, and thus exclude access of individual tropocollagen molecules (diameter of ∼15 Å), not to mention collagen assemblies. To confirm this on a biochemical level for bacterial collagenases, we investigated the temperature dependence of tropocollagen and gelatin degradation using the CU of ColQ1 from *Bacillus cereus strain Q1* (ColQ1-CU) and found tropocollagen degradation was more sensitive to decreasing the reaction temperature than gelatin degradation ^37^ (**Fig. S1**). These results emphasize that the unwinding of collagen is an essential prerequisite for collagen cleavage by bacterial collagenases.

### Recognition of the hierarchical substrate collagen by clostridial collagenases

Physiological processing of collagen involves three distinct recognition events, binding to (i) insoluble fibrillar collagen; (ii) soluble tropocollagen; and (iii) unfolded α-chains (*aka* gelatin). Therefore, we systematically examined the affinity of full-length ColG, its CU and of its individual domains for these three hierarchical levels of collagen. Protein variants that contained the PD carried the inactivating E524A mutation. Full-length ColG E524A (Y119-K1118) (ColG-FL E524A) and ColG-AD (Y119-S392) contained an N- or C-terminal maltose-binding protein tag, respectively.

First, we investigated the binding of ColG-FL E524A and of its domains to insoluble collagen fibers, the predominant physiological substrate (**Fig. 2a**). We observed concentration-dependent binding to fibers of ColG-FL E524A, ColG-CBD2 (N1004-K1118), ColG-CU E524A (Y119-G790), and ColG-AD in decreasing order, whereas very low or no binding to insoluble collagen was seen for ColG-PD E524A (D398-G790) and ColG-PKD (I799-N880). Of all ColG-domains including the CU, ColG-CBD2 bound best to collagen fibers, reflecting the eponymous ‘collagen-binding domain’ function, while the PD showed no detectable binding. It is, therefore, remarkable that ColG-CU E524A showed a higher affinity to collagen fibers than the single AD, indicating a non-additive, but cooperative binding mode of AD and PD to the fibrillar substrate. Together, these results suggest that the highly affine binding of the full-length enzyme is based on the high affinity of the CBD and the CU to collagen fibers.

**Fig. 2:**
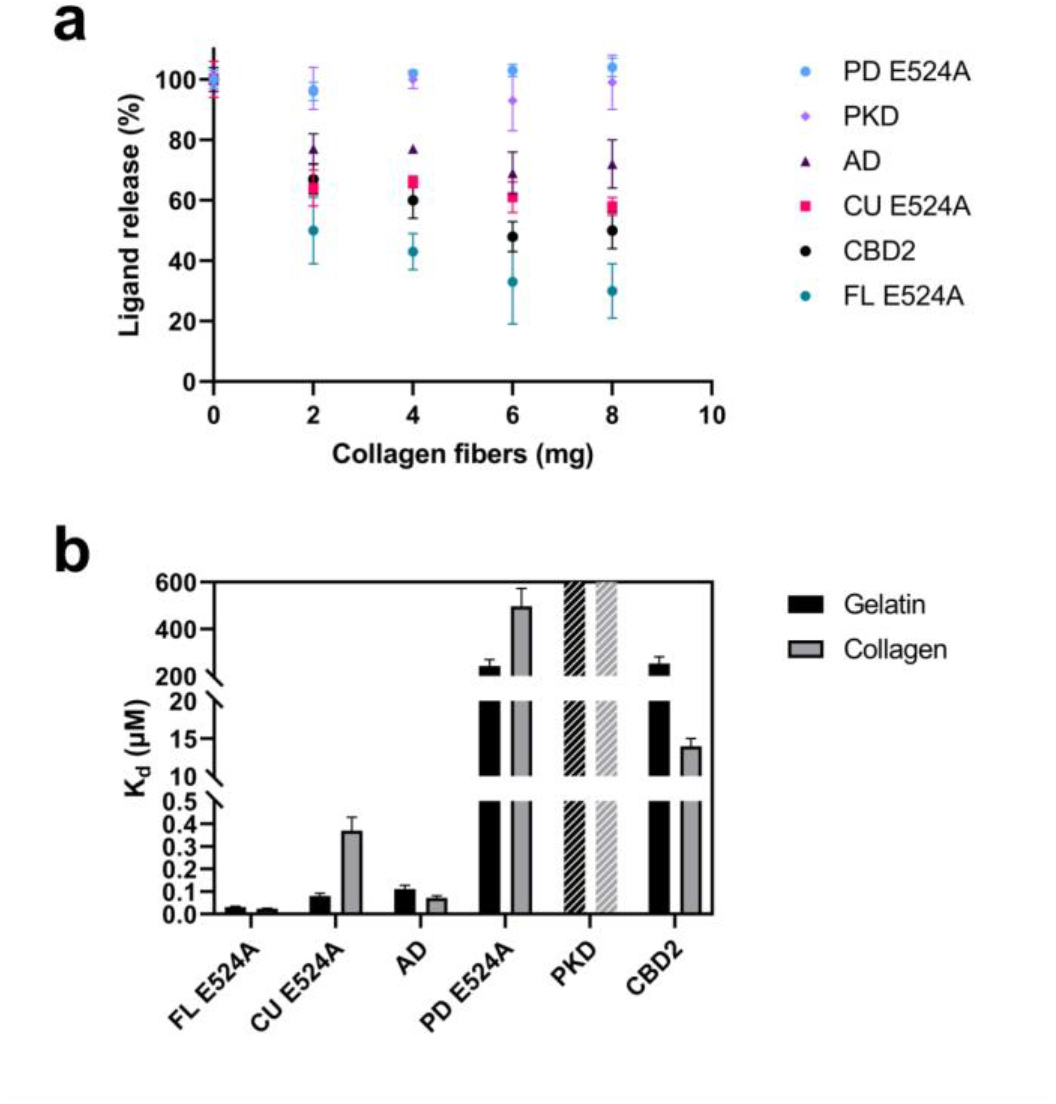
Binding of ColG variants to insoluble collagen fibers, soluble collagen and gelatin. **a**, Release of fluorescently labelled ColG variants from insoluble collagen fibers. **b**, Binding affinities to soluble type I collagen and type I gelatin. Apparent dissociation constants were determined via indirect ELISA, whereby the apparent K_d_ values represent the binding to a gelatin or collagen substrate each with multiple binding sites per molecule. The binding curves are shown in Fig. S1.

Second, we examined the binding towards tropocollagen and gelatin (**Fig. 2b**). Among all variants, ColG-FL E524A displayed the highest affinity to both substrates. Intriguingly, it displayed very similar affinities towards unwound and folded collagen (apparent dissociation constant (K_d_) = 0.031 ± 0.004 µM *vs*. 0.024 ± 0.002 µM, respectively). Most of its affinity towards soluble collagen and gelatin was mediated by the CU. ColG-CU E254A bound best to gelatin (Kd = 0.08 ± 0.01 µM), and showed an ∼ 5-fold lower affinity to collagen (Kd = 0.37 ± 0.06 µM). Interestingly, the PD and accessory domains showed mostly poor binding towards gelatin and collagen, except for CBD2. ColG-CBD2 bound to triple-helical collagen with a Kd of 14 ± 1 µM, but it displayed low binding affinity to gelatin (Kd = 253 ± 30 µM). In contrast, ColG-PKD exhibited very little binding to both substrates; the binding was so poor that no Kd values could be determined, but the data suggest a Kd of well over 1 mM for both substrates (**Fig. S2**).

ColG-FL displayed a more than twofold higher affinity towards gelatin and a ∼9-fold higher affinity to collagen compared to ColG-CU. These increases in affinity result most likely from the C-terminal CBDs. In fact, the differential CBD binding affinities to collagen and gelatin consistently explain the preferential collagen binding of ColG-FL, compensating for the weaker collagen binding of ColG-CU (**Fig. 2b**).

To dissect CU binding to gelatin and collagen, we analyzed the binding of the individual AD and PD, which together form the CU. ColG-AD exhibited a Kd of 0.11 ± 0.02 µM for gelatin and an even better Kd of 0.072 ± 0.009 µM for collagen. In contrast, ColG-PD E524A showed only low binding to gelatin (Kd of 243 ± 28 µM) and a twofold lower binding towards collagen (Kd of 497 ± 76 µM). When comparing the affinities to gelatin and collagen of the AD and the PD with the CU, first, it is striking that the CU binds to gelatin like the AD (Kd = 0.08 ± 0.01 µM *vs*. 0.11 ± 0.02 µM). This suggests that binding to gelatin is predominantly mediated via the AD, consistent with the several hundred-fold difference in binding affinities of AD and PD. Second and counterintuitively, CU binding to triple-helical collagen (Kd = 0.37 ± 0.06 µM) cannot be accounted for by an additive effect of the AD (Kd = 0.072 ± 0.009 µM) and the PD (Kd = 497 ± 76 µM), but rather suggests an antagonistic interaction of both when binding to the triple-helical substrate.

### Bacterial collagenases unwind collagen only locally, not globally

To investigate the triple-helicase activity of ColG, we measured the melting temperature (T_m_) of tropocollagen in the absence and presence of inactive ColG-CU G494V by circular dichroism spectroscopy (CD). α-Chymotrypsin served as negative control (**Fig. 3**). We observed no significant change in the T_m_, neither in the absence or the presence of ColG-CU G494V, nor of α-chymotrypsin (**Fig. 3a**). This suggests that ColG does not globally unwind triple-helical collagen, but that the unwinding only takes place locally. Next, we performed a proteolytic complementation assay using ColQ1-PD as ‘cutter’ enzyme and the inactive ColQ1-CU G472V as ‘triple helicase’ (**Fig. 3b**, **Fig. S3, Fig. S4)**. Despite the addition of increasing concentrations of ColQ1-CU G472V to a constant high amount of ColQ1-PD, we observed no turnover of natively folded collagen, suggesting that the unwinding of collagen by ColQ1-CU G472V only occurred locally and reversibly.

**Fig. 3:**
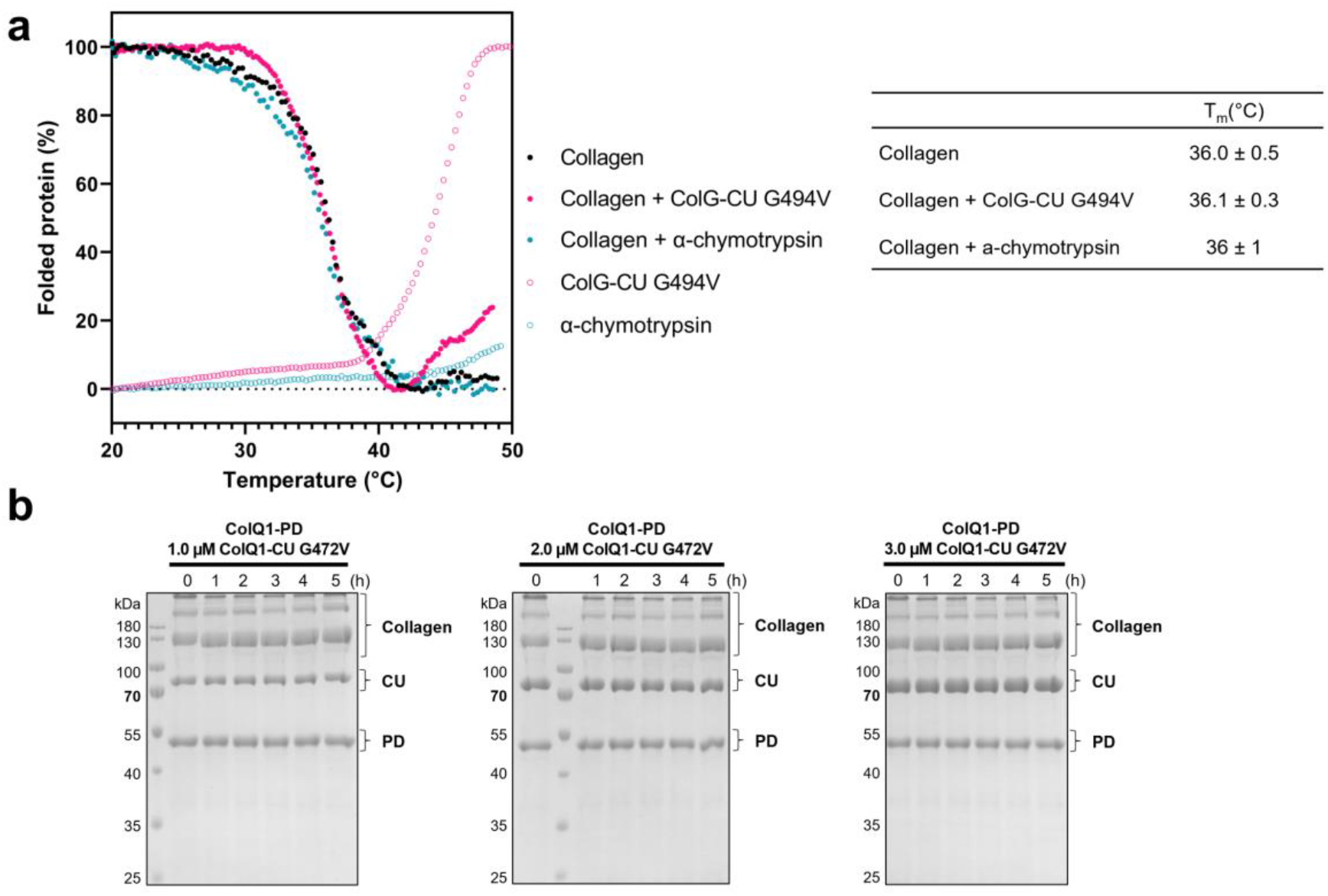
ColG does not unwind type I collagen globally. **a**, Thermal transition curves and melting temperatures from CD experiments for type I collagen with or without ColG-CU G494V and α-chymotrypsin. **b**, SDS-PAGE analysis of a degradation complementation assay, in which collagen was incubated with equimolar concentrations of ColQ1-PD and increasing concentrations of ColQ1-CU G472V at 25 °C. The integrity of the substrate was verified by co-incubation with 0.83 μM α-chymotrypsin (data not shown).

### Unwinding of short collagen triple helices can be monitored via CD

The local unwinding of the collagen triple helix is not accessible to detection by CD. We concluded that the collagen fraction that is unwound by the collagenase is too small compared to the dimensions of the whole molecule (∼300 nm in length, ∼338 Gly-X-Y tripeptides/α-chain) to have a significant effect on the overall T_m_ of tropocollagen. Therefore, we generated a short collagen-like substrate V-(GXY)_26_ to monitor the unwinding activity. This construct is derived from prokaryotic Scl2 (*Streptococcus pyogenes* collagen-like protein 2). It is composed of the trimerization V-domain of Scl2.3 and the proline-rich B-segment of the CL domain of Scl2.28 ^38,39^. The native B-segment forms a stable collagen-like triple-helix with a T_m_ of 29.5 °C ^39^. We modified segment B to increase ColG’s affinity by mutating four of its tripeptides to Gly-Pro-Ala (**Fig. S5**) ^31^. This modified construct V-(GXY)_26_ was efficiently recognized by ColG-CU. The triple-helical B-segment was readily degraded by ColG-CU, leaving the V-domain trimer intact, while the inactive G494V mutant could not cleave it. α-chymotrypsin could only partially cleave the V-domain, but it left the triple helix intact (**Fig. 4a**).

**Fig. 4:**
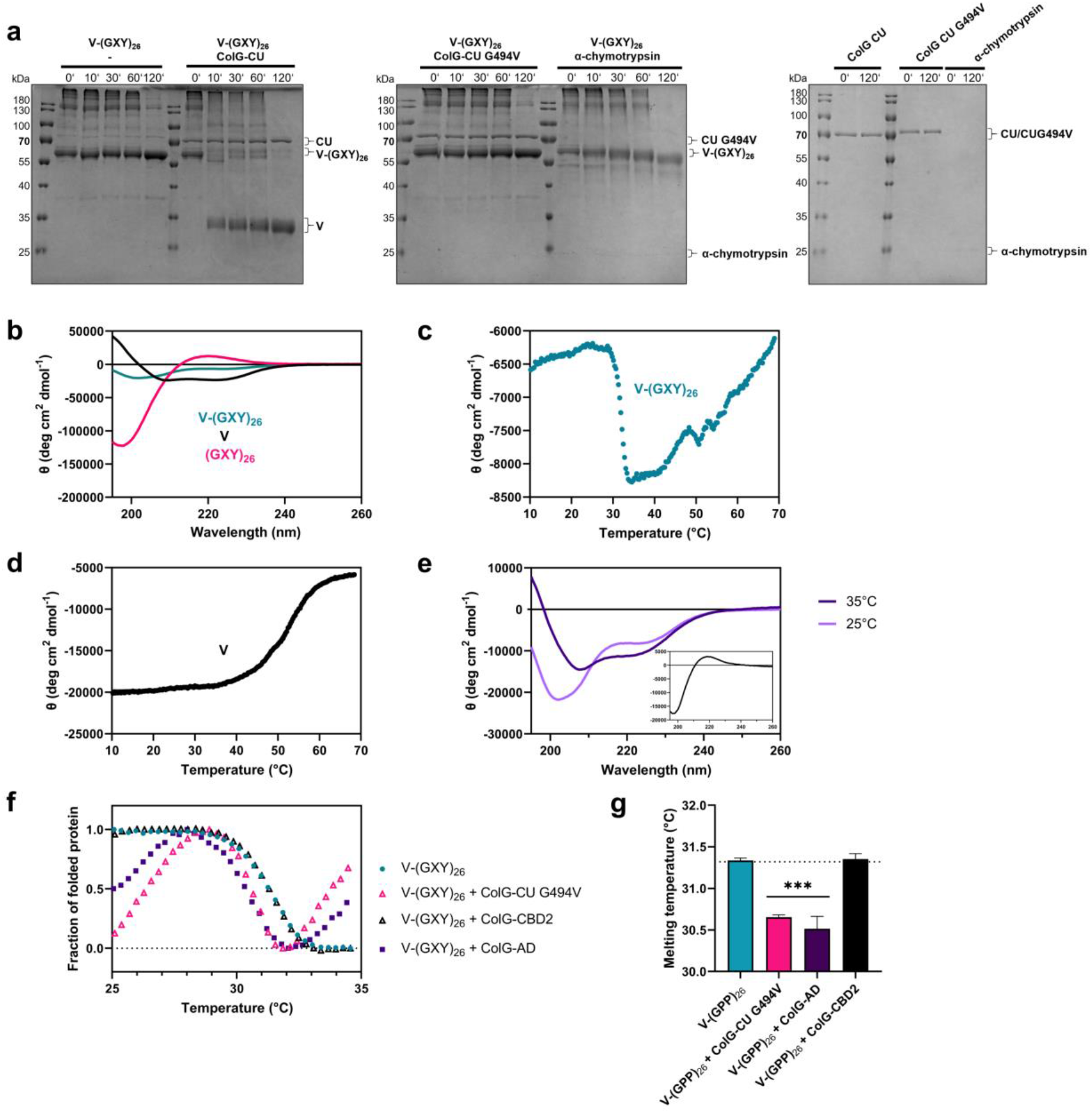
Triple-helicase activity of ColG-CU is detectable via CD spectroscopy. **a-b**, V-(GXY)_26_ is a short collagen-like substrate. **a**, SDS-PAGE analysis of time course of incubation of V-(GXY)_26_ with and without ColG-CU WT, ColG-CU G494 and α-chymotrypsin. Note that the V-domain migrates as an SDS-stable trimer at an apparent mass of 30 kDa; similarly, V-(GXY)_26_ migrates as a trimer. **b**, CD spectrum of V-(GXY)_26_ and of (GXY)_26_, prepared by pepsin digest of V_Scl2.3_-(GXY)_26_. **c-d**, Melting profiles of V-(GXY)_26_ (**c**) and of the single V-domain (**d**) determined by CD spectroscopy. **e**, CD spectra of V-(GXY)_26_ recorded at 25 °C and 35°C. The inlet shows the CD difference spectrum of the 25 °C spectrum minus the 35°C spectrum. **f**, Melting profiles of the short collagen mimic V-(GXY)_26_ in absence or presence of ColG-CU G494V, ColG-AD and ColG-CBD2 monitored via CD spectroscopy. **g**, Melting temperatures of V-(GXY)_26_ in absence or presence of ColG-CU G494V, ColG-AD and ColG-CBD2 (mean ± SD).

We confirmed the triple-helical fold of the B-segment of V-(GXY)_26_ using CD (**Fig. 4b,e**). The net spectrum of V-(GXY)_26_ is composed of the α-helical contribution of the V-domain (78 aa) and the triple-helical contribution of segment B (78 aa). The isolated modified B-segment, *i.e.,* (GXY)_26_, displayed the spectral fingerprint of triple-helical collagen with a negative band below 200 nm and a positive band at 222 nm. When monitoring the molar ellipticity at 222 nm at temperatures between 10 and 70 °C, the melting of the V-domain (T_m_ = 52 ± 2 °C) could be clearly differentiated from the melting of the triple-helical (GXY)_26_ part (T_m_ = 31.82 ± 0.05 °C) (**Fig. 4c-d**). Moreover, the difference spectrum of V-(GXY)_26_ from spectra recorded at 25 °C and 35 °C, *i.e.*, before and after B melting, showed CD transitions typical for the collagen triple helix ^40^ (**Fig. 4e**).

### The activator domain is a triple helicase

To identify the triple-helicase entity in ColG, we tested all ColG domains which we had found to bind to triple-helical collagen (ColG-CU, ColG-AD and ColG-CBD2) for their effect on the denaturation of V-(GXY)_26_. Remarkably, in the presence of a twofold excess of ColG-CU G494V a distinct destabilization of the triple helix was observed, causing a decrease in T_m_ from 31.34 ± 0.02 °C to 30.66 ± 0.02 °C (**Fig. 4f-g**). As a control we added a 6-fold excess of ColG-CBD2 to V-(GXY)_26_, which is known to bind to triple-helical collagen, but which cannot unwind collagen ^23^. The presence of ColG-CBD2 did not alter the T_m_ of V-(GXY)_26_ (31.35 ± 0.05 °C). To our knowledge, this is the first time that the triple-helicase activity of a collagenase could be directly demonstrated.

Given that triple-helical collagen preferentially binds to the AD (**Fig. 2b**), we measured the melting profile of V-(GXY)_26_ in presence of ColG-AD and found a clear decrease in T_m_ of V-(GXY)_26_ to 30.52 ± 0.15 °C, comparable to the temperature shift detected in presence of ColG-CU G494V (**Fig. 4f-g**). This finding suggests that the AD can unfold a triple-helix independently of the PD.

To confirm the triple-helicase activity of the AD, we performed two substrate degradation assays (**Fig. 5**). First, we monitored the cleavage of soluble collagen using ColQ1, which cleaves folded collagen six times more efficiently than ColG ^37^ (**Fig. 5a**, **Fig. S6a-d**). Yet, co-incubation of tropocollagen with increasing concentrations of ColQ1-AD in the presence of a constant amount of ColQ1-PD resulted only in minute degradation fragments. We hypothesized that the apparent failure of the two domains to functionally complement each other for collagen degradation was caused by the large substrate dimensions, resulting in a vast binding surface compared to the small number of enzyme molecules added.

**Fig. 5:**
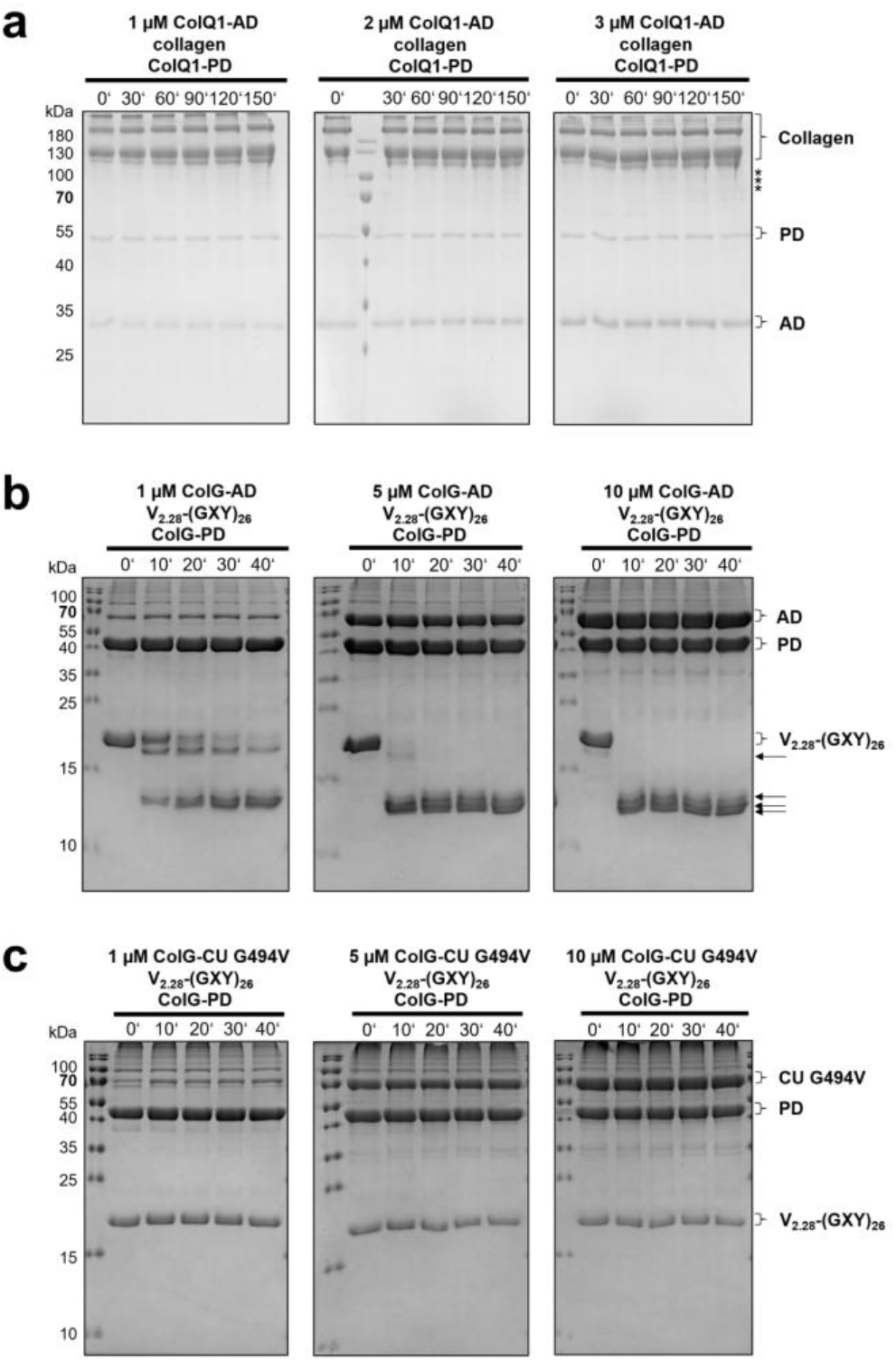
Local and transient triple-helicase activity of AD requiring simultaneous co-localization of the PD for collagen cleavage. **a**, SDS-PAGE analysis of time course of co-incubation of 3.33 µM collagen with increasing concentrations of ColQ1-AD (1-3 µM) in the presence of 0.6 µM ColQ1-PD at 25 °C. Minor fragments are highlighted with an asterisk. **b-c**, SDS-PAGE analysis of time course of co-incubation of 10 µM V_2.28_-(GXY)_26_, which migrates as monomer, with 1, 5 or 10 µM ColG-AD-MBP (**b**) or 1, 5 or 10 µM ColG-CU G494V (**c**) in the presence of 10 µM ColG-PD. Control reactions can be found in **Fig. S8**.

When we repeated the assay using a smaller triple-helical substrate instead of tropocollagen, *i.e.*, V_2.28_-(GXY)_26_, a modified version of V-(GXY)_26_ containing the V-domain of Scl2.28 ^39^, we observed a clear dose-dependent degradation of V_2.28_-(GXY)_26_, which depended on the simultaneous presence of ColG-PD as ‘cutter’ and of ColG-AD as ‘triple helicase’ (**Fig. 5b**, **Fig. S7a-b**). The higher local concentrations of the single AD and PD on the surface of the model collagen allowed for the productive interaction of both for the degradation of the triple-helical substrate. These results show that the activator domain on its own is able to functionally unwind triple-helical collagen, *i.e*., the AD is the minimal entity enabling peptidolytic collagen degradation.

### AD and PD must simultaneously colocalize for efficient catalysis of triple-helical collagen

Inspired by complementation experiments by the Nagase group, we next tested whether a CU-dead mutant could complement an active gelatinase to degrade triple-helical collagen, as was successfully shown for MMP-1 ^36^. Therefore, we repeated the degradation assay with V_2.28_-(GXY)_26_ as substrate shown in **Fig. 5b**, but replaced ColG-AD as ‘unwinder’ by ColG-CU G494V (**Fig. 5c**). Interestingly, in presence of ColG-CU G494V the isolated ColG-PD could no longer process the collagen mimic, indicating that the unwound peptide chain(s) were not accessible to the single PD when ColG-CU G494V was bound to V_2.28_-(GXY)_26_. Consequently, the unwinding must be short-lived and reversible, as sequential unwinding by the CU-dead mutant followed by gelatinase cleavage was unproductive.

### AD residues critical for collagen binding and unwinding

To identify AD residues crucial for substrate recognition and unwinding, we performed a bioinformatical analysis of M9B collagenases, followed by an alanine scan of strictly/highly conserved surface-exposed residues on the inner AD surface based on the crystal structure of ColG as reference model (PDB: 2y6i). Seven candidate residues were identified (ColG: F148, E191, R194, Y198, Y201, N251, and F295). We generated the homologous single-point mutants of ColQ1-CU and its inactive variants ColQ1-CU G472V (ColQ1: F123, E166, R169, Y173, F176, N226, and Y270) to investigate their effects on binding to gelatin and tropocollagen, triple-helix unwinding, and hydrolysis. The structural integrity of single-point mutants was confirmed for active variants using a peptidolytic assay (**Fig. S9**), and for inactive variants by nanoDSF (**Fig. S10**). In addition, the proper folding of inactive variants was confirmed via CD spectra taken before melting experiments.

Intriguingly, all seven mutants displayed impaired tropocollagen turnover *in vitro*, while they were able to cleave peptides like the WT (**Fig. 6**, **Fig. S9**). In contrast, three control mutants (ColQ1: Y251A, N317A, Y321A, also surface-exposed AD residues) did not show any aberrant collagen turnover (**Fig. S11**).

**Fig. 6:**
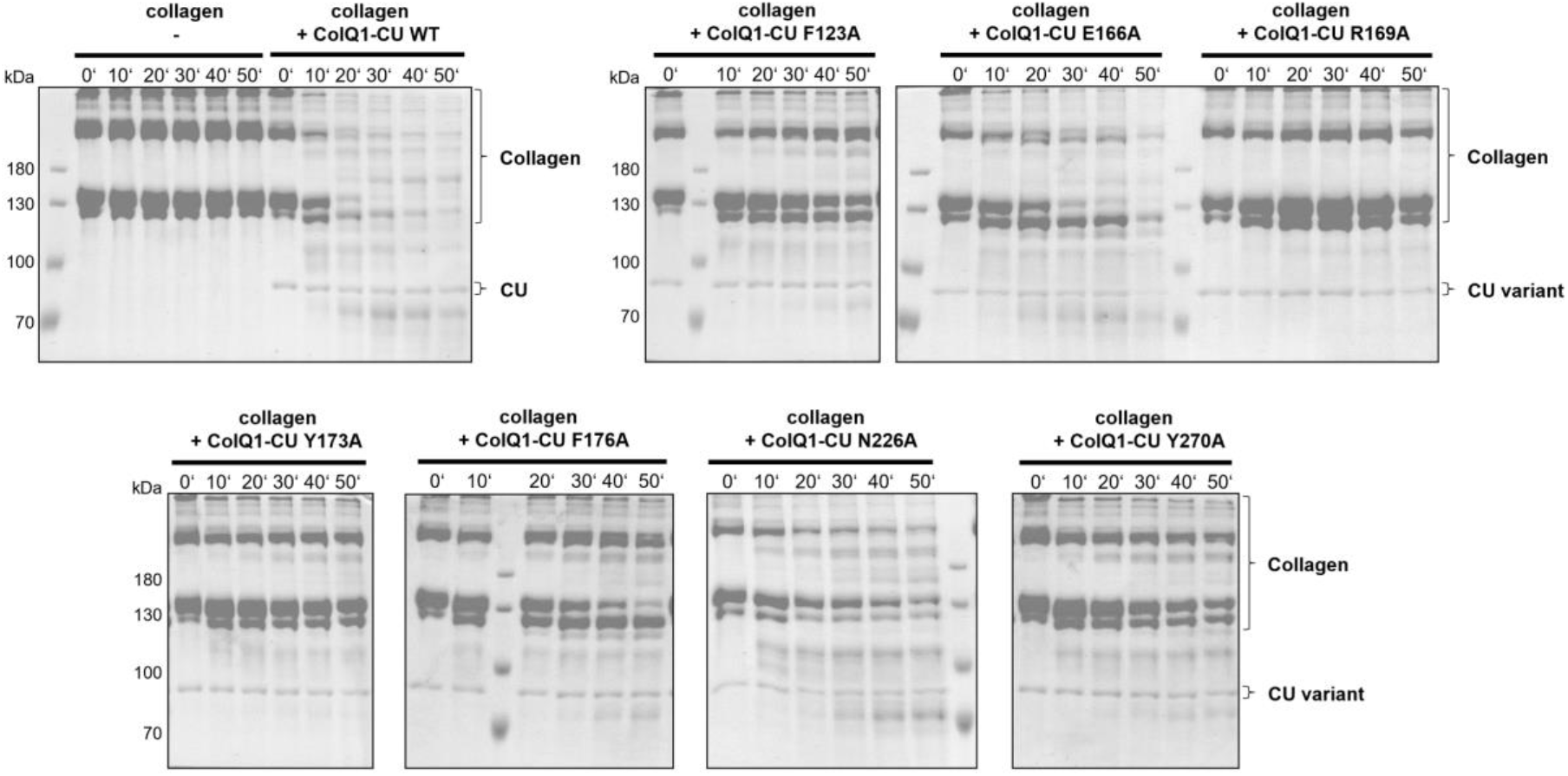
Activity of ColQ1-CU WT and its mutants towards soluble type I collagen. SDS-PAGE analysis of time course of 3.33 µM type I collagen co-incubation with 0.2 µM ColQ1-CU variants at 25 °C.

To determine whether this deficiency in collagen degradation was the result of binding and/or unwinding defects, we investigated (i) the binding affinities towards gelatin and soluble collagen using ELISA assays, and (ii) the effects on the triple-helicase activity when co-incubated with the CD reporter substrate V-(GXY)_26_ (**Fig. 7 & Table S1**).

**Fig. 7:**
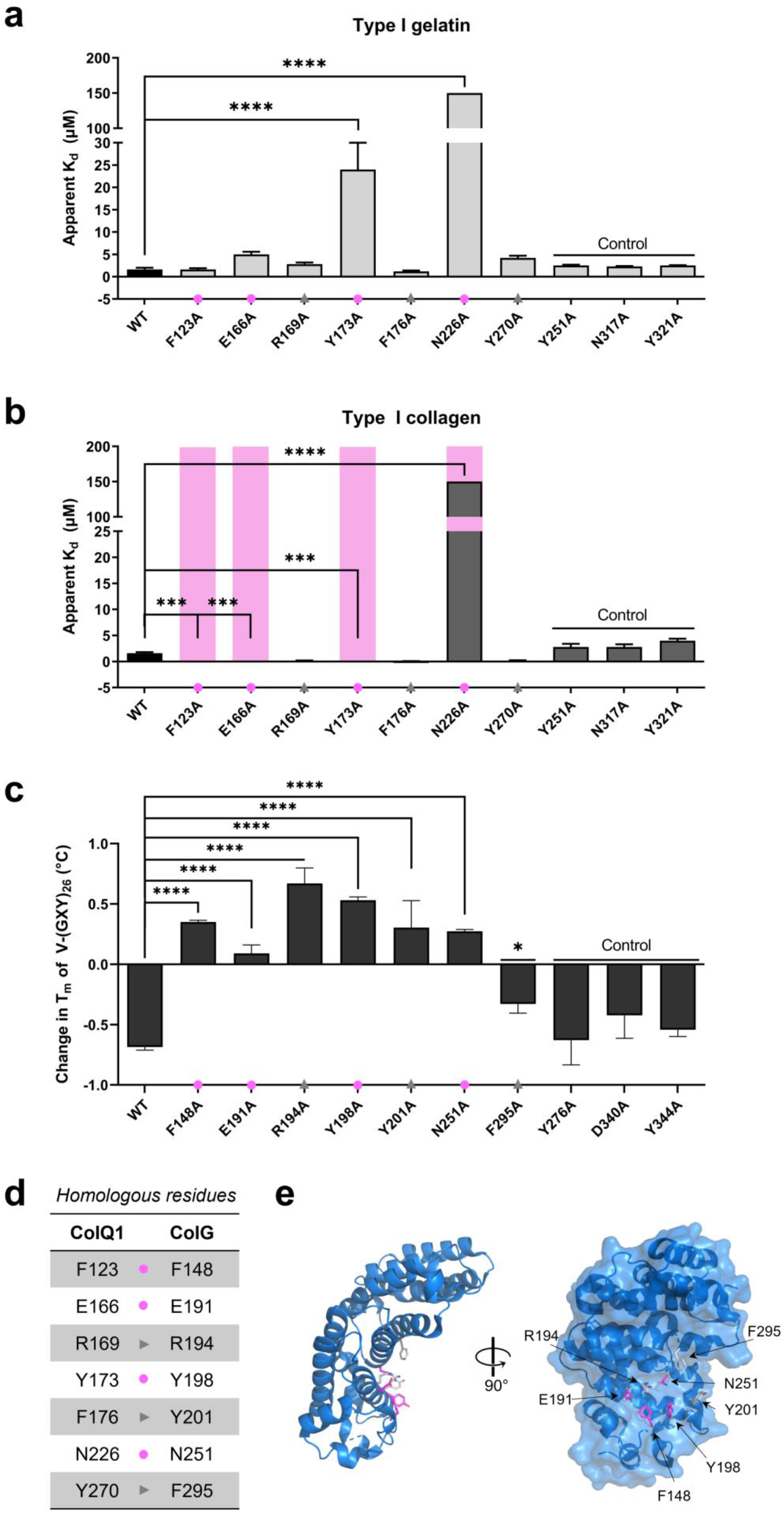
Effect of single-point mutations in AD on binding affinities towards type I gelatin and soluble type I collagen, and on triple-helicase activity. **a**, K_d_ values of ColQ1-CU G472V (=WT) and mutants were determined by indirect ELISA (**a**-**b**). The binding affinities towards collagen were tested at 25 °C to preserve the triple-helical fold, while the binding affinities towards gelatin were tested at 37 °C. Loss of affinity was marked with a magenta bar if no binding at 100 µM concentration was detected. **c**, Melting temperatures of V-(GXY)_26_ in presence of ColG-CU G494V (= WT) or its mutants determined by CD spectroscopy. Homologous variants of ColQ1 and ColG are arranged on top of each other (**a-c**) in the figure. **d**, Table of the homologous residues in ColQ1-AD and ColG-AD that were investigated. **e**, Ribbon and surface representation of the AD. Residues important for collagen binding, i.e., F148, E191, Y198, and N251 (magenta), and the residues crucial for unwinding, i.e., R194, Y201, and F295 (grey), are highlighted as sticks. To ease comparison between the panels, the residues involved in collagen binding (magenta circle) and unwinding (grey triangle) are marked.

We compared the binding of seven AD mutants to the WT (**Fig. 7a-b**). ColQ1-CU G472V (WT) bound similarly to gelatin and soluble collagen (K_d_ = 1.6 ± 0.4 µM vs. 1.6 ± 0.2 µM). Two residues crucial for gelatin binding in ColQ1-AD were identified, Y173 and N226. The binding affinities to gelatin dropped drastically – 15-fold and 90-fold, respectively – when these residues were replaced by alanine. Both residues were also important for collagen binding. In total, four mutations resulted in significant binding defects to soluble collagen, *i.e.* F123A, E166A, Y173A and N226A. Similarly to gelatin binding, the N226A mutation resulted in a more than 90-fold reduction in binding affinity to collagen. Even more crucial were the mutations F123A, E166A and Y173A, which caused a complete loss of binding.

These four residues critical for collagen binding and degradation cluster in the central AD cavity (**Fig. 7e**), forming a prominent exosite. In striking contrast, three other functionally inactivating mutations R169A, F176A and Y270A resulted in a marked increase in affinity of the CU to collagen of 9-fold, 20-fold, and 8-fold, respectively. To understand this, we examined the effect of AD mutations on the triple-helicase activity using the homologous mutations in inactive ColG-CU G494V (**Fig. 7c-d**). The four AD mutations, which had resulted in binding defects to collagen in ColQ1-CU (ColG-F148A/ColQ1-F123A, ColG-E191A/ColQ1-E166A, ColG-Y198A/ColQ1-Y173A, and ColG-N251A/ColQ1-N226A), also failed to unwind V-(GXY)_26_. This result confirms that triple-helix binding is a prerequisite for unwinding. Interestingly, ColG-R194A/ColQ1-R169A and ColG-Y201A/ColQ1-F176A, which had displayed a higher affinity to soluble collagen than the WT, were also not able to lower the T_m_ of V-(GXY)_26_ upon co-incubation. This suggests that these two residues are critical for unwinding. To a lesser degree, this was also true for ColG-F295A/ColQ1-Y270A. This finding indicates that triple-helix unwinding is not only dependent on integral binding properties, but it relies also on specific interactions which interfere directly or indirectly with the triple-helical structure, *e.g.,* by disturbing the local hydration shell.

### Conformational trapping of the CU through intramolecular crosslinking

To investigate the effect of the relative AD-PD geometry and dynamics in more detail, we generated conformationally trapped variants of ColG-CU^41–43^. Here, conformational trapping of ColG-CU was achieved by either engineering intramolecular disulfides that crosslink the AD and the PD at various positions or by deletion of the AD-PD linker. The positions of the crosslinks were spread along the CU axis with the aim to trap the CU in a semi-closed *vs*. closed conformation (**Fig. 8a-b**). We briefly summarize the design, production and quality assessment of crosslinked variants generated on the basis of a cysteine-free reference CF (C218S/C262S) in the supplement ^44^. We investigated three conformationally-restrained variants, two disulfide-linked variants, ColG-CU CL3 (E294C/T483C) and CL4 (C prior to N-terminus/D586C), and a Δlinker variant (ΔG389-V397). The activity towards short peptides was unaffected in these three conformationally restrained and the CF variants ^23,44^.

**Fig. 8:**
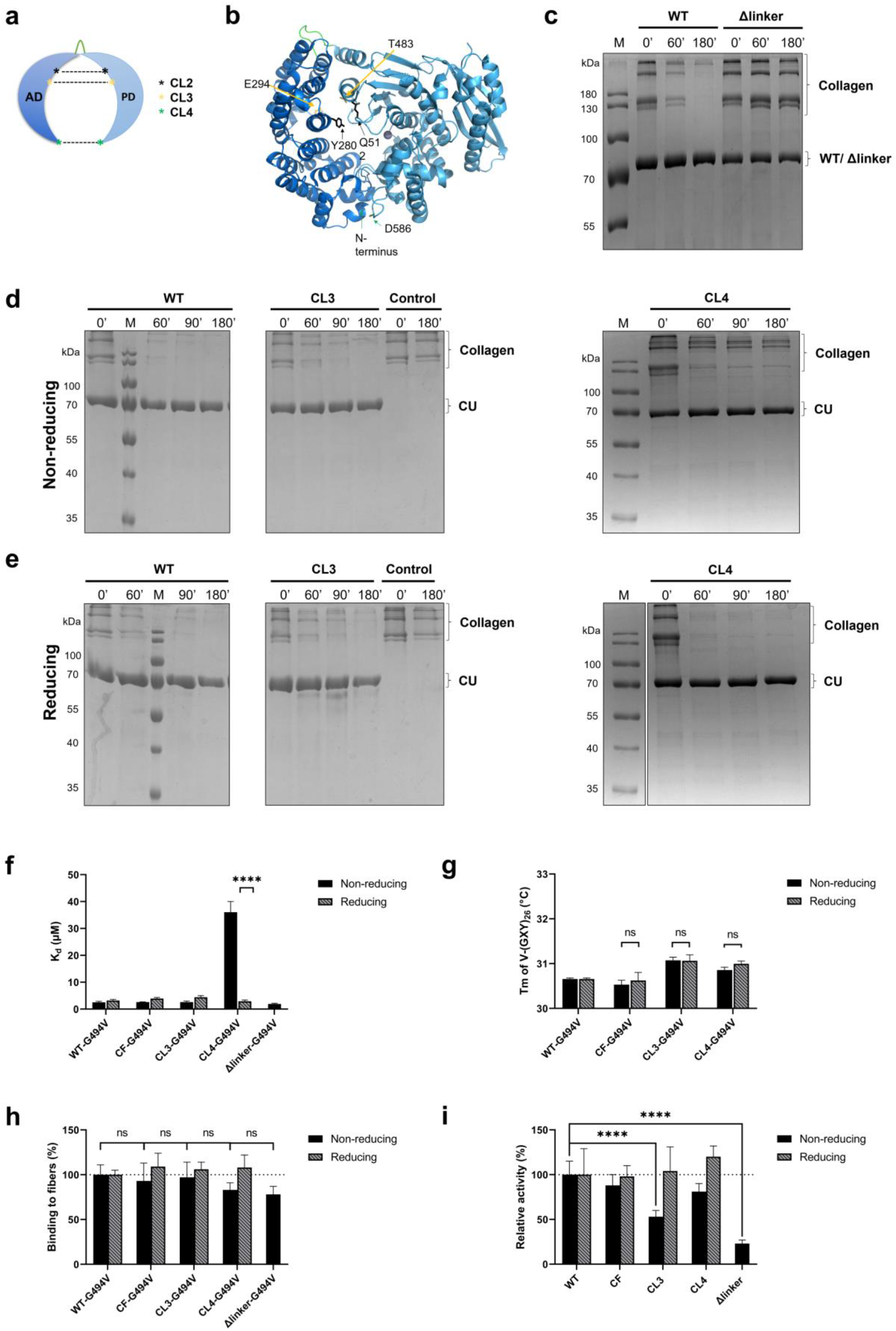
Conformational trapping and its effect on triple-helix degradation and binding. **a**, Scheme of crosslinked mutants. **b**, Model of (semi)-closed conformation of ColG-CU. Mutation sites for the introduction of cysteines are shown in sticks. **c-e,** SDS-PAGE analysis of time course of degradation of soluble type I collagen by ColG-CU variants in absence (**c**, **d**) or presence (**e**) of 10 mM ß-mercaptoethanol. **f,** Binding affinity of ColG-CU G494V (=WT G494V) and the mutants were determined by microscale thermophoresis. The binding curves can be found in **Fig. S15**. **g,** Melting profiles of the collagen mimic V-(GXY)_26_ in presence of WT G494V and the mutants determined by CD spectroscopy. Experiments (**f**, **g**) were performed in presence or absence of 1 mM ß-mercaptoethanol. **h,** Binding to insoluble collagen fibers monitored via ligand release assay using fluorescently labelled, inactive ColG-CU. **i,** Degradation of insoluble collagen fibers variants monitored via fluorescamine-citrate assay. Experiments (**h**,**i**) were performed in presence or absence of 10 mM ß-mercaptoethanol.

### Triple-helical collagen turnover, but not binding is significantly reduced in the open conformation of the CU

In CL3, the engineered crosslink is located ∼8 Å above the level of the active-site cleft. It is designed to trap the CU in a semi-closed conformation. In CL4, where the AD and PD are linked at their tips, this will maintain a closed conformation (**Fig. 8b**); whereas the linker deletion (Δlinker) enforces a quasi-open CU conformation.

Looking at the activity of Δlinker against tropocollagen compared to the WT, we observed a drastic inhibitory effect. The linker deletion nearly abolished collagen degradation (**Fig. 8c**). This result suggests that in the open conformation cleavage of the triple helix is severely hampered, most likely because substrate presentation to the PD is compromised. This conclusion is supported by the observation that the binding affinity towards tropocollagen was unchanged compared to the WT (**Fig. 8f**).

When we compared the collagenolytic activities of CL3 and CL4 in the crosslinked vs. open state, the results were remarkable (**Fig. 8d-e**). The crosslink in CL3 did not compromise tropocollagen turnover, and no significant difference between crosslinked and non-crosslinked state was observed, suggesting that the semi-closed conformation did not interfere with binding and processing of the triple helix. Only the conformationally most restricted variant CL4 showed a 6-fold slower collagen turnover than the WT. This reduction in collagen turnover could be completely reversed by opening of the crosslink (**Fig. 8d-e**).

To see whether this decrease in tropocollagen cleavage was the result of impaired triple-helix binding and/or compromised helix unwinding, we examined the binding and unwinding behavior of the crosslinked mutants (CL3, CL4; CF served as control; all building on the inactivate G494V mutation) and compared them to the reduced mutants and ColG-CU WT. We found no significant change in binding affinity to V-(GXY)_26_ in CF-G494V and CL3-G494V compared to WT-G494V, neither under non-reducing conditions, nor under reducing conditions (**Fig. 8f**). Remarkably, however, we observed a ∼12-fold drop in binding affinity of CL4-G494V in the crosslinked state compared to the non-crosslinked state (K_d_ = 36 ± 4 µM *vs*. 2.9 ± 0.4 µM). The closed conformation led to a sharp decrease in binding affinity, explaining the observed decrease in tropocollagen turnover by CL4 in the non-reduced state.

We examined also the triple-helicase activity of the crosslinked CU variants, however, we found no ß-mercaptoethanol-dependent effect on the melting temperatures of V-(GXY)_26_ in the presence of CL3-G494V and CL4-G494V (**Fig. 8g**).

### Conformational trapping impairs cleavage of collagen fibers, but not their binding

Finally, we investigated the effect of conformational trapping of ColG-CU on the recognition and processing of insoluble collagen fibers (**Fig. 8h-i**). Importantly, we observed no significant binding defects in all mutants compared to WT-G494V, with an only marginally lower binding in crosslinked CL4-G494V and Δlinker-G494V. Thus, the closed conformation did not result in a major decrease in binding affinity. Apparently, in the disulfide-restrained CU synergistic AD-PD interactions with collagen fibrils were able to compensate for effects of impaired cooperativity between the AD and the PD in the context of triple-helical collagen (**Fig. 8h**, **Fig. 2a**).

Regarding the processing of collagen fibers by the mutants, the findings were remarkable. Δlinker with its locked-open conformation showed a drastic reduction of 77 ±4% in collagen-fiber cleavage compared to the WT (**Fig. 8i**), consistent with a similarly reduced cleavage of soluble collagen (**Fig. 8c**). CL3, trapped in a semi-closed conformation, showed also a notable 47 ± 7% decrease in fiber processing, which could be completely reversed by the addition of ß-mercaptoethanol. Finally, CL4 exhibited a 19 ± 9% reduction in activity in the crosslinked state, and full activity could be recovered upon disulfide reduction, as in CL3. All three conformationally restrained variants demonstrate the necessity of inter-domain dynamics for fibrillar collagen degradation, but not binding.

## Discussion

Collagen is hierarchically organized, with soluble triple-helical collagen assembling into insoluble microfibrils and fibers. The data presented here show that bacterial collagenases, by virtue of their multi-domain structures, recognize, disassemble and process these different collagen conformers in distinct and previously underappreciated ways.

Our data reveal that the CBD2 of ColG binds fibrillar collagen more tightly than the AD, while the binding affinities are reversed for soluble collagen and gelatin; this indicates that CBD2 employs multiple binding sites in collagen fibers. These multivalent interactions cannot be explained by avidity effects only, which should also apply to the AD. Possibly, additional binding motifs such as pyridinoline crosslinks, which are specific to supramolecular fibers, selectively modify the binding affinities of CBD2 and AD. As suggested by Caviness *et al*.^45^, in ColG CBD2 is likely to initiate binding to fibrillar collagen. Both CBDs present in ColG work then in tandem to anchor the full-length enzyme to the fibril, facilitating its movement toward the C-terminus of the fibril.

The AD dominates the binding interaction toward triple-helical collagen, whereas the isolated PD shows no or only negligible binding affinity to fibrillar and soluble collagen, respectively. It was, therefore, unexpected to see that the binding-incompetent PD significantly impacts the binding to collagen in the context of the CU. Even more surprising, the effect had an opposite sign for soluble-and fibrillar-collagen binding. While the PD interacts synergistically with the AD to increase the binding affinity of the CU to insoluble collagen fibers (**Fig. 2a**), the opposite is true for soluble collagen (**Fig. 2b**), where the PD antagonizes AD binding. These apparently counterintuitive observations must be rationalized mostly by entropic considerations, as the PD’s enthalpic contributions to binding are negligible. The binding affinity is determined by the ratio of the on-rate (k_on_) and the off-rate (k_off_). The presence of the PD abates the diffusion-controlled accessibility of a large rigid macromolecular ligand such as soluble or fibrillar collagen, but not that of small or unfolded flexible macromolecular ligands such as gelatin, where the locally controlled diffusion experiences no steric restrictions to access the AD ^46^. Only suitably pre-oriented triple helices will be able to access the saddle-shaped topology of the CU and bind to the AD. This conformational selection reduces the k_on_, and it represents a significant entropic cost for the binding of soluble and fibrillar collagen. This effect is further enhanced when considering that AD binding to collagen is most likely a multi-step process, where an initial docking geometry is subsequently adjusted for optimal interaction. The presence of the PD sterically interferes with this multi-step docking process. To avoid misconceptions, we wish to make it clear that the CU will be significantly more mobile than collagen, reflecting their different hydrodynamic radii. However, since only relative movements are relevant in the docking process and it is more conventional to assume that the substrate moves and binds to the enzyme receptor, we adhere to this linguistic convention.

However, the situation differs for the off-rate in case of soluble and (micro)fibrillar collagen (**Fig. 9**). In case of the AD-bound triple helix, the k_off_ to dissociate off and leave the AD cavity is hardly reduced by the presence of the PD. The diffusion distances to leave the CU saddle are comparable to those, where collisions with the PD could significantly hinder and slow down soluble collagen to diffuse out of the CU saddle. Taken both effects together, the affinity (*i.e.*, the ratio k_on_ / k_off_) of CU towards soluble collagen is reduced. By contrast, the saddle-shaped topology of the CU acts as a cage for bound (micro)fibrillar collagen. Most diffusion events away from the AD collide with the PD, thus slowing down k_off_ significantly, more than k_on_ is reduced by the presence of the PD. The selective effect of the CU topology on k_off_, and the overall reduction of k_on_ thus explain the observed antagonistic effect towards conformationally rigid triple helices and the synergistic effect towards fibrillar collagen (**Fig. 9**).

**Fig. 9:**
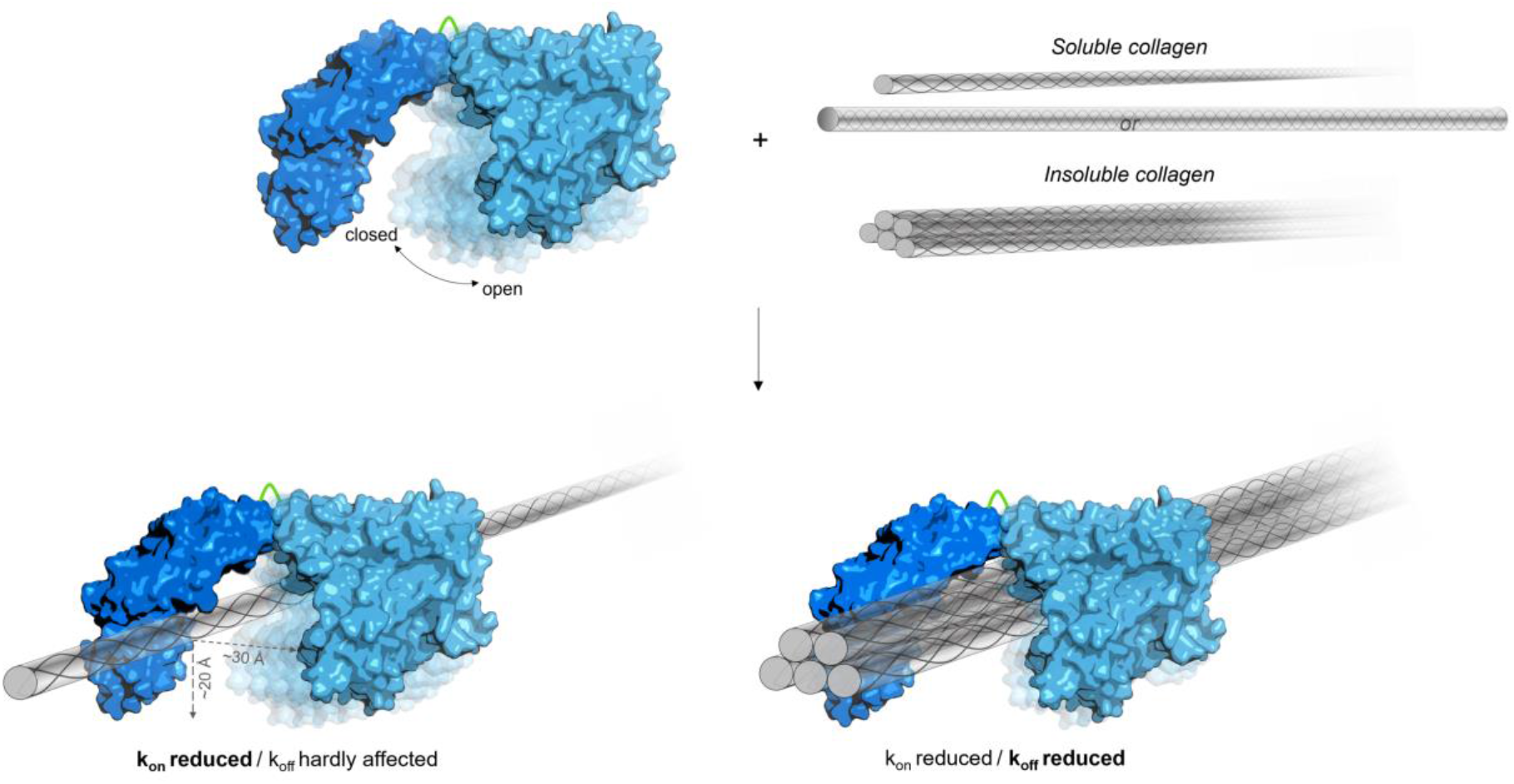
Schematic model of CU binding to soluble and insoluble collagen illustrating antagonistic and synergistic effects. The presence of the PD sterically restricts access to AD binding sites in the saddle-shaped CU for soluble collagen molecules and microfibrils. Only suitably pre-oriented molecules can access and bind, resulting in an overall reduction of k_on_. Compared to the AD, the affinity of the CU towards soluble collagen is, however, lowered, because the presence of the PD hardly diminshes k_off_ for the relatively small triple helix, whereas the saddle-shaped CU can ‘cage’ the considerably larger microfibril, resulting in a substantial k_off_-effect and thus in an overall higher affinity of the CU for microfibrils.

AD’s prominent role is not limited to collagen binding. We show here that the AD can unwind collagen triple helices. Monitoring AD-induced unfolding of collagen is challenging because the unfolding is locally restricted, transient and reversible (**Fig. 3-5**), but became feasible by developing a mini-collagen reporter for CD spectroscopy. Moreover, we identified and characterized individual AD sites, which inactivated its unwinding activity either by loss of collagen binding or by gain of collagen binding and triple-helix stabilization (**Fig. 6**, **Fig. 7**). Specifically, we identified four residues in the AD that are crucial for the recognition of triple-helical collagen (ColG-F148/ColQ1-F123, ColG-E191/ColQ1-E166, ColG-Y198/ColQ1-Y173, and ColG-N251/ColQ1-N226), of which ColG-Y198/ColQ1-Y173 and ColG-N251/ColQ1-N226 are also key residues for binding to unfolded collagen. Importantly, however, we identified three further AD residues, which did not interfere with collagen binding, but proved vital for the unwinding of collagen. When mutated to alanine (ColG-R194A/ColQ1-R169A, ColG-Y201A/ColQ1-F176A, and ColG-F295A/ColQ1-Y270A), collagenolytic activity was abolished. Wang *et al.* identified F107, R153, and Y157 as important residues for triple-helix binding in the VhaC-AD.^29^ We found ColG-F148/ColQ1-F123 and ColG-Y198/ColQ1-Y173, which are homologs of VhaC-F107 and VhaC-Y157, also to be vital for collagen binding: Interestingly, however, the mutation ColG-R194A/ColQ1-R169A behaved contrary to VhaC-R153A. While VhaC-R153A resulted in a ∼70% reduced binding affinity to collagen, ColQ1-R169A displayed a nearly 10-fold higher affinity towards the triple-helical substrate compared to ColQ1-WT. Moreover, all three residues with improved collagen binding (*i.e.,* ColG-R194A/ColQ1-R169A plus ColG-Y201A/ColQ1-F176A, and ColG-F295A/ColQ1-Y270A) were essential for triple-helix unwinding. Thus, next to the lack of a noticeable linker between the AD and PD, and of a catalytic helper domain^29^, this supports the idea that there are subtle differences in the way collagen is processed by M9A and M9B collagenases, respectively.

Finally, we tested whether the discovered synergistic and antagonistic domain interactions were consistent with the proposed chew-and-digest model ^23^. To this end, we used conformationally restrained CU variants, which either locked the CU in an open state by minimizing the linker between AD and PD or by disulfide linking the AD with the PD, thus locking in more closed CU conformations (**Fig. 8a-b**). Overall, the relative flexibility of the AD-PD domains was critical for processing of soluble and fibrillar collagen substrates, as the degradation of these collagens was largely abolished by the locked-open Δlinker variant, although its binding was unaffected. Mechanistic details of collagen processing depend on the collagen conformer serving as substrate. When analyzing the disulfide-locked variants, the reduced AD-PD interdomain flexibility affected the processing of soluble collagen less severely than that of fibrillar collagen (**Fig. 8**). These findings can be explained by a modified chew-and-digest mechanism of bacterial collagenolysis. The Δlinker and the disulfide crosslinked variant CL4 restrain the CU in two opposing conformations, *i.e.,* open and closed, respectively. The locked-closed conformation (crosslinked CL4) keeps the collagenase “mouth” closed, thereby restraining the chewing amplitude, but not preventing substrate chewing altogether. Low amplitude chewing allows for crunching of relatively small substrates (soluble collagen), but much less so of larger substrates (fibrillar collagen), consistent with our findings. By contrast, in the locked-open conformation (Δlinker), chewing is impossible and so is processing. Although intuitively plausible, why is collagen processing impossible in the locked-open collagenase? After all, substrate binding to the AD is unaffected, and AD binding is sufficient to induce collagen unwinding (**Fig. 4f**). So whenever unwound collagen dissociates from the AD, the PD should be able to cleave the prepared, gelatin-like substrate. However, this was not observed, because of the transient and reversible nature of the collagen unwinding. As soon as the locally unwound collagen dissociates from the AD, it instantaneously refolds to its triple helical conformation. For productive cleavage, it is necessary to deliver the AD-bound collagen to the active site of the PD. This conclusion is consistent with the results of the complementation experiment, where an AD, but not an inactive CU, could feed unfolded collagen productively to an active PD (**Fig. 5**). The inactive CU locally unfolds the collagen but at the same time protects it from cleavage, because its inactive PD shields the unfolded substrate.

This finding also demonstrates that the binding and substrate presentation of triple-helical collagen decisively differ between collagenolytic MMPs and bacterial collagenases, while they share some basic features. In both, the catalytic domains require the help of an additional activating domain to be able to process triple-helical collagen. In both families, the peptidase domains alone possess little affinity for folded collagen ^35^. And in both enzyme families, collagen unwinding is locally restricted ^36^, yet distinctly different. In MMP-1, the catalytic and hemopexin domains bind to the triple helix in an elongated conformation, which leaves the surface of the bound substrate largely exposed ^47^. Consistent with this finding, the single catalytic domain of MMP-1 added as “cutter” enzyme could cleave triple-helical collagen when pre-incubated and unwound by the inactivate full-length MMP-1 E200A ^36^. The same is, however, not true for bacterial collagenases. Here, access to the unwound α-chain(s) is blocked in the CU, because the triple helix is sandwiched between the AD and the PD. This contracted conformation is required for proper presentation of the α-chain to the PD for cleavage.

## Conclusion

Within multi-domain enzymes, the catalytic domain is usually considered to be the most important domain because it contains the active site, where substrate processing takes place. Our data presented here provide a much more resolved picture of substrate recognition and processing in collagenases, reflecting the complexity of collagen substrates. The AD initiates collagen binding (together with the CBDs), unfolds collagen by destabilizing its triple-helical structure, and presents the locally unwound collagen to the active site of the PD. While substantial cooperation between the AD and PD was to be expected, it was as a complete surprise to see synergistic and antagonistic cooperativity for fibrillar and soluble collagen, respectively. The here obtained deeper mechanistic understanding of substrate selectivity paves the way for the design of collagenases with unprecedented specificities. Such uniquely specific enzymes can be used for various biotechnological applications in histology ^48^, wound healing ^49^, or oncolysis ^50^, to name a few.

## Materials and methods

### Materials

Acid-soluble type I atelocollagen from bovine hides was purchased from Cell Guidance Systems (UK). Type I gelatin was produced by heating the acid-soluble type I atelocollagen for 5 min to 95°C. Type I collagen fibers from bovine Achilles tendon were purchased from Merck (Germany).

### *In silico* analysis of the AD in bacterial collagenases

Multiple-sequence alignment of the AD of ColA from *Clostridium perfringens* (P43153), ColG from *C. botulinum* (B2TJU5), ColT from *C. tetani* (Q899Y1), ColG and ColH from *H. histolyticum* (Q9X721 & Q46085), ColQ1 from *Bacillus cereus* (B9J3S4) were performed using Clustal Omega ^51^.

### Construction of V-(GXY)_26_

The coding sequences of V-B from Scl2.28 incorporating the 4 mutated tripeptides and the coding sequence of the V domain from Scl2.3 were purchased from Genscript (Germany). The modified coding sequence of V-B was cloned into a modified pET15b vector. The endogenous V domain was replaced by the V domain from Scl2.3 via Gibson assembly. All constructs were verified by sequencing at Eurofins Genomics (Germany).

### Construction of ColG-CU variants

Based on a pET15b expression plasmid of ColG-CU WT (Tyr119–Gly790) ^23^, a plasmid encoding a cysteine-free ColG-CU variant (CF) (C218S, C262S) was generated by Gibson assembly. Using site-directed mutagenesis via inverse PCR ^52^, the plasmids for the ColG variants CL1, CL2, CL3, and CL4 were obtained from the cysteine-free template, while the linker variants were generated from the plasmid encoding ColG-CU WT. All constructs were verified by sequencing at Eurofins Genomics (Germany).

### Expression and purification of ColG variants, ColQ variants and V-(GXY)_26_

All protein variants were expressed in *E. coli* Nico21 (DE3) cells and purified as described previously ^37,53^. Monodisperse protein fractions were collected and stored at -80 °C, and the sample purity was confirmed by SDS-PAGE analysis under denaturing conditions.

### Preparation of (GXY)_26_

11 µM V-(GXY)_26_ were co-incubated in a molar ratio of 1:20 with pepsin in 50 mM acetic acid pH 3.0 at 4 °C for 72 h. The reaction was stopped by pH adjustment to pH 7.4 and the addition of 10 mM ß-mercaptoethanol. The structural integrity of (GXY)_26_ was confirmed by co-incubation with α-chymotrypsin in a molar ratio 1:10 for up to 3 h (data not shown).

### Disulfide formation and thiol quantification assay

For the crosslinkage, purified collagenases (0.1 mg/ml final concentration) were suspended in 50 mM Tris-HCl pH 8.5, 1 mM β-mercaptoethanol, 300 mM NaCl, 5% glycerol, 1 mM CaCl_2_ and 3 mM NaN_3_. The oxidation reactions were kept at 4 °C for 10 days followed by centrifugation (16,500 g for 20 min). The non-oxidized molecules were removed by Activated Thiol Sepharose^TM^ 4B chromatography (Sigma Aldrich) performed according to the manufacturer’s recommendations. In short, the clarified samples were loaded onto a self-packed activated thiol sepharose 4B column equilibrated in 50 mM Tris-HCl pH 8.5, 300 mM NaCl, 5% glycerol, 1 mM CaCl_2_ and 3 mM NaN_3_ and incubated overnight at 4 °C. The crosslinked variants were collected in the flowthrough fraction. The crosslinked monomers were separated from misoxidized aggregates by size exclusion chromatography using a Superdex 200 10/300 GL (Cytiva, Germany) and 10 mM HEPES pH 7.5, 100 mM NaCl, 5% glycerol and 3 mM NaN_3_ as buffer. The extent of disulfide-bridge formation in the samples CL1-CL4 was examined using the thiol-specific fluorochrome 7-diethylamino-3-(4-maleinimidophenyl)-4-methyl coumarin (CPM) ^54^ and ColG-CU WT (contains 2 cysteines) as positive control. CPM (Sigma-Aldrich, Germany) was dissolved at 4 mg/ml in DMSO and stored at -80 °C. Prior to use, the stock was diluted in the reaction buffer (10 mM HEPES pH 7.5 and 100 mM NaCl supplemented with 20% DMSO). The assay was performed in a total volume of 120 µL. The protein samples were diluted in the reaction buffer to 1.0 µM. 10 µL diluted dye and 16 µL DMSO were added to 94 µL protein solution. After 3 min of incubation at 60 °C for protein denaturation, the fluorescence was measured in an Infinite M200 plate reader (Tecan, Austria) at an excitation and emission wavelength of 387 nm and 463 nm, respectively.

### Circular dichroism spectroscopy

Far UV circular dichroism (CD) spectra in the wavelength range from 195 to 260 nm were recorded using a Chirascan Plus CD Spectrophotometer (Applied Photophysics, Leatherhead, UK) at 25 °C. The instrument was flushed with nitrogen, the pathlength was 0.5 mm, spectral bandwidth was set to 1 nm and the scan time per point 1 s. The samples (2.25 µM type I collagen, 5.0 µM V-(GXY)_26_, 5.0 µM V, 2.25 µM and 10.0 µM ColG-CU G494V, 30.0 µM ColG-CBD2, 2.25 µM α-chymotrypsin, 10 µM ColG-CU G494V mutants, 10.0 µM CF, and 10.0 µM CL1-CL4) were measured in 15 mM Tris-SO_4_ pH 7.5, 100 mM NaF, 1 mM CaCl_2_ and ± 1 mM ß-mercaptoethanol. Melting profiles were measured at similar concentrations at a wavelength of 222 nm from 10 °C to 70 °C or 20 °C to 50 °C (setting time: 120 s, scan time per point: 1 s, step: 0.3 °C) and from 25 °C to 35 °C (setting time: 240 s, scan time per point: 10 s). To verify the proper folding of the samples, CD spectra from 195 to 260 nm were recorded at the start temperature prior to initiating the temperature ramp. In case of the melting temperatures of V-(GXY)_26_ in absence or presence of ColG-CU G494V, ColG-AD and ColG-CBD2, the statistical significance was determined by one-way ANOVA followed by Dunnett’s test for multiple comparisons (***P < 0.001). When comparing the melting temperatures of the V-(GXY)_26_ in the presence of the different crosslinked variants, the statistical significance was determined by one-way ANOVA followed by Holm-Šídák’s multiple comparisons test, comparing the reduced variant to its non-reduced control.

### Determination of thermal stability

Thermal denaturation assays were performed using Tycho NT. 6 (NanoTemper Technologies, Germany). The measurements were performed at 0.1 mg/ml protein concentration in 15 mM Tris-SO_4_ pH 7.5, 100 mM NaF and 1 mM CaCl_2_ in triplicates. Intrinsic fluorescence was recorded at 330 and 350 nm while heating the sample from 35 to 95 °C at a rate of 30 °C/min. Fluorescence ratio (350/330 nm) and inflection temperature were calculated by Tycho NT. 6 software.

### Peptidolytic assay

The peptide-degradation assay was performed as described previously ^55^. In short, all ColG-CU variants were tested at a final concentration of 16 nM and co-incubated with 2 µM of the quenched-fluorescent peptidic substrate Mca-Ala-Gly-Pro-Pro-Gly-Pro-Dpa-Gly-Arg-NH2 (FS1-1) (Mca = (7-Methoxycoumarin-4-yl)acetyl; Dpa = N-3-(2,4-dinitrophenyl)-L-2,3-diaminopropionyl). The final reaction buffer contained 250 mM HEPES pH 7.5, 400 mM NaCl, 10 mM CaCl_2,_ 10 µM ZnCl_2_, 2% DMSO and 3 mM NaN_3_. Reactions were performed in the presence and absence of 0.5 mM TCEP. Cleavage of the substrate was monitored for 2 min at 25 °C (excitation: 328 nm, emission: 392 nm) in an Infinite M200 plate reader (Tecan, Austria) and the initial velocity (v_0_) was determined from the progress curves (<10% substrate conversion).

Steady state measurements were used to determine K_M_ for FS1-1. The final concentration of ColQ1-CU WT and its variants was 1 nM and the concentration of the substrate was screened from 0 – 175 µM. The initial velocity was determined from the progress curves (<10% substrate conversion) using linear regression and inner filter effect-correction ^56^. The ε_ex328_ and ε_em392_ were estimated in the buffer used for the kinetic assay to be 21794 M^-1^ cm^-1^ and 9561 M^-1^ cm^-1^. K_M_ was calculated by non-linear regression from the resulting Michaelis − Menten plot using GraphPad Prism 9.1.2 (Graph Pad Software, San Diego, CA, USA).

### Degradation of V-(GXY)_26_ monitored via SDS–PAGE

10 µM V-(GXY)_26_ were digested at 25 °C by 0.5 µM ColG-CU or 0.5 µM α-chymotrypsin in 250 mM HEPES pH 7.5, 400 mM NaCl, 10 mM CaCl_2_, and 10 µM ZnCl_2_ for up to 2 h at 25 °C. Samples were taken at indicated time points and the reaction was stopped by addition of SDS-PAGE loading buffer on ice. The degradation was monitored using 12% non-reducing SDS-PAGE gels.

### Degradation of V_2.28_-(GXY)_26_ monitored via SDS–PAGE

10 µM V_2.28_-(GXY)_26_ were co-incubated with 1, 5 or 10 µM ColG-AD-MBP in the presence or absence of 10 µM ColG-PD in 250 mM HEPES pH 7.5, 400 mM NaCl, 10 mM CaCl_2_, and 10 µM ZnCl_2_ for up to 40 min at 25 °C. Samples were taken at indicated time points and the reaction was stopped by addition of SDS-PAGE loading buffer on ice. The degradation was monitored using 16% non-reducing SDS-PAGE gels.

### Degradation of soluble collagen monitored via SDS–PAGE

1 mg/ml acid-soluble type I atelocollagen from bovine hides (Cell Guidance Systems, Cambridge, UK) was digested at 25 °C by 4.54 µM collagenase in 250 mM HEPES pH 7.5, 400 mM NaCl, 10 mM CaCl_2_, 10 µM ZnCl_2_, and ± 10 mM ß-mercaptoethanol for up to 4 h. Samples were taken at indicated time points and the reaction was stopped by addition of 38 mM EDTA. The integrity of the triple-helical collagen fold was verified by co-incubation with 0.83 µM α-chymotrypsin (FLUKA, Switzerland) (data not shown). The degradation was monitored on 12% non-reducing SDS-PAGE gels. All experiments were performed at least in triplicates. Densitometric analysis were performed using GelAnalyzer 19.1 (www.gelanalyzer.com).

### Collagen-fibril degradation monitored via fluorescamine-citrate assay

Two milligrams of insoluble fibrillar collagen from bovine Achilles tendon (Merck, Germany) were added to an Nanosep microcentrifugal device (0.2 µm pore size) with a low-binding Bio-Inert membrane (Pall, Germany). 500 µL reaction buffer (250 mM HEPES pH 7.5, 400 mM NaCl, 10 mM CaCl_2_, and 10 µM ZnCl_2_, ± 10 mM ß-mercaptoethanol) were added for 15 min at room temperature to swell the fibers and then removed via centrifugation (13,000 g for 2 min). Then 200 µL 1.0 µM ColG-CU variants were added and incubated for 2 h at room temperature. The filtrate was collected and supplemented with 38 mM EDTA to stop the reaction. The amount of hydrolysis was quantified in comparison to the control reaction with ColG-CU WT exploiting the N-terminal-specific adduction of fluorescamine to peptides proteins at mildly acidic pH ^57^. In short, 5 µL of the stopped reaction were diluted 1:10 with reaction buffer and then mixed 1:1 with 1 M citrate pH 5.6. 100 µL of this mixture were added to 10 µL 2.5 mg/ml fluorescamine in acetone and incubated for 5 min at room temperature, before the fluorescence was measured at 25 °C (excitation: 390 nm, emission: 475 nm) in an Infinite M200 plate reader (Tecan, Austria). All experiments were performed at least in triplicates.

### Indirect ELISA

Since gelatin can partially refold into triple-helical structures upon cooling ^40–42^, particular care was taken to use gelatin concentrations and reaction temperatures that disfavored triple-helix formation during coating and binding assays. For the coating, 1 mg/ml bovine type I atelocollagen stock solutions were prepared in 0.1 M HCl and diluted to 5 µg/ml in 20 mM phosphate buffer pH 7.4, 150 mM NaCl. Gelatin solutions were prepared by heating type I collagen stock for 5 min at 95 °C. 96-well high-binding microplates (Greiner Bio-One, Germany) were incubated with 100 µL coating solution per well overnight at 4 °C for collagen plates, and at 37 °C for gelatin plates. After incubation, the plates were washed four times with PBST. Coated wells were blocked with 1x PBS supplemented with 10% skim-milk for 90 min and then washed four times with PBST. Collagen plates were stabilized with PBST supplemented with 1% BSA fraction V and 5% sucrose. Plates were dried and stored at 4 °C. For the binding assay, the hexahistidine-tagged ColG variants were prepared in PBST supplemented with 1% BSA fraction V. For K_D_ determination, the samples were serially diluted 1:3. 75 µL sample per well were incubated for 140 min with collagen at room temperature, while gelatin plates were incubated at 37 °C for the same time period. The plates were washed four times with PBST and were then incubated with 1:15,000 rabbit polyclonal 6x His-tag antibody conjugated to HRP (Abcam, Austria) for 1 h at room temperature, followed by four washes with PBST and a final wash with 1x PBS. As substrate 75 µL 3,3′,5,5′-tetramethylbenzidine were added per well. The peroxidase activity was followed by measuring the absorption at 650 nm every 15 s for 225 s in an Infinite M200 plate reader (Tecan, Austria) at 25 °C. The initial velocity was determined by regression analysis. For K_d_ determination, the data was fitted to the Hill equation using GraphPad Prism 9 (Graph Pad Software, USA). The apparent dissociation constant K_d_ is given as mean values of three independent experiments ± standard deviation. Statistical significance was determined by one-way ANOVA followed by Dunnett’s test for multiple comparisons (* P < 0.05, *** P < 0.001, **** P < 0.0001).

### Binding assay to insoluble collagen fibers

ColG-variants were labeled using the Monolith NT™ Protein Labeling Kit RED-NHS 2nd Generation Amine reactive (NanoTemper, Germany) and the yield of labelled protein and the degree of labelling were determined according to the manufacturer’s manual. Insoluble type I collagen from bovine Achilles tendon (Sigma, Germany) (0, 2, 4, 6, and 8 mg or 6 mg) was prewetted with 225 µL reaction buffer (50 mM Hepes pH 7.5, 100 mM NaCl, 10 mM CaCl_2_, 0.1% Tween-20, 1% fraction V of bovine serum albumin, 3 mM NaN_3_, ± 10 mM ß-mercaptoethanol) for 15 min at RT, before 100 µL labelled ColG-variants solubilized in reaction buffer were added. The final concentration of labelled protein was 0.2 µM, except for ColG-PKD, which was added at 5 µM final concentration to ensure a proper signal-to-noise ratio. The reactions were incubated at 25 °C for 30 min with stirring in the dark and then centrifuged at 13,000 g for 5 min at RT. Labelled ColG variants in reaction buffer without any substrate were used as controls. The collagen-binding ability of labelled proteins was determined monitoring the free fluorescence intensity in the supernatant after incubation using a Tecan M200 Infinite plate reader (Tecan, Austria) (647 nm excitation/680 nm emission).

### Binding assay to V-(GXY)_26_ monitored via microscale thermophoresis

Microscale thermophoresis experiments were performed on a NanoTemper Monolith NT.115 instrument (NanoTemper Technologies). ColG variants were labeled using the Monolith NT™ Protein Labeling Kit RED-NHS 2nd Generation Amine reactive (NanoTemper Technologies, Germany). After labeling, the ColG variants were eluted into 250 mM Hepes pH 7.5, 150 mM NaCl, 10 mM CaCl_2_, 10% glycerol, 3 mM NaN_3_ and stored at -80 °C. The yield of labelled protein and the degree of labelling were determined according to the manufacturer’s manual. For the assay, the labelled proteins and the ligand were diluted into 50 mM Hepes pH 7.5, 150 mM NaCl, 10 mM CaCl_2_, 10 µM ZnCl_2_, 0.05% Tween-20, 3 mM NaN_3_, ± 1 mM ß-mercaptoethanol. The ColG variants were used at a concentration of final concentration of 25 nM (except for 9xAla-Linker where 40 nM were used because of the low degree of fluorescent labelling), while the ligand was titrated in a 1:2 dilution series. After that ColG variants and ligand were mixed 1:2 and the samples were centrifuged for 10 min at 17,000 g at RT. The solutions were immediately transferred into Monolith NT.115 standard capillaries and measured using 60% excitation power at 22 °C. The binding of the ligand caused a reduction in the initial fluorescence signal, confirmed via specificity test performed according to the manufacturer’s guidelines. The experiments were performed in triplicates. The change in the initial fluorescence signal was used to calculate the apparent binding constant K_d_ using non-linear regression analysis in GraphPad Prism 9 (Graph Pad Software, USA).

### Statistics and reproducibility

Data analyses were carried out using GraphPad Prism 9 (Graph Pad Software, USA). All data are shown as means ± standard deviations from three independent experiments. For CD spectroscopy analysis, similar results were obtained from three independent experiments. Detailed data analyses are described in the text.

## Acknowledgments

We thank C. Cabrele for valuable discussions during project design.The authors are grateful for funding by the Austrian Science Fund (FWF), P31843.

## Author contributions

J.S. performed experiments (cloning, protein production, quality control of crosslinked variants, binding and enzymological experiments), analysed data and wrote the draft; A.C.W. performed experiments (cloning, protein production, CD experiments) and analysed data; A.H. performed experiments (protein production), H.B. devised the project and wrote the paper, E.S. devised the project, performed experiments (cloning, protein production, binding and enzymological experiments, CD experiments), analysed the data, prepared the manuscript and the figures, and wrote the paper.

