## Supplementary_Material for "The activator domain of bacterial collagenases drives collagen recognition, unwinding and processing"

**Tables of contents**

1. **Supporting figures and tables**
2. **Experimental section**
3. **References**

### 1. Supporting figures and tables

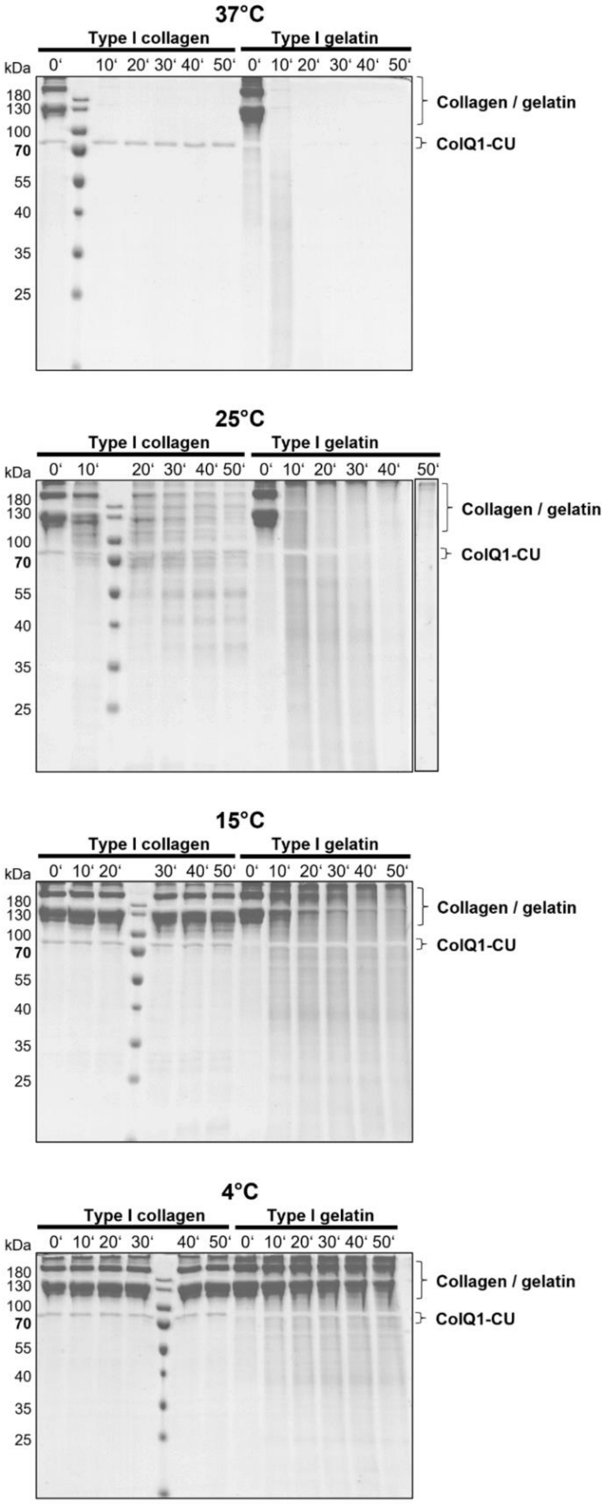


**Fig. S1: Temperature dependence of collagen and gelatin degradation by ColQ1-CU.** SDS-PAGE analysis of the time course of soluble type I collagen and type I gelatin degradation, respectively, by ColQ1-CU performed at 37 °C, 25 °C, 15 °C and 4 °C, respectively. At 37 °C and 25 °C, tropocollagen and gelatin were readily. At 15 °C, collagen turnover nearly halted, while gelatin degradation continued, yet at a significantly slower rate. At 4 °C, no degradation of collagen could be observed, but cleavage of gelatin still proceeded at a very slow rate. Thus, whereas gelatin cleavage was qualitatively governed by the Arrhenius equation[1,2], tropocollagen degradation was more sensitive to decreasing the reaction temperature, indicative for the existence of an additional distinct transition temperature for collagen as compared to gelatin.

**a** **Type I gelatin**

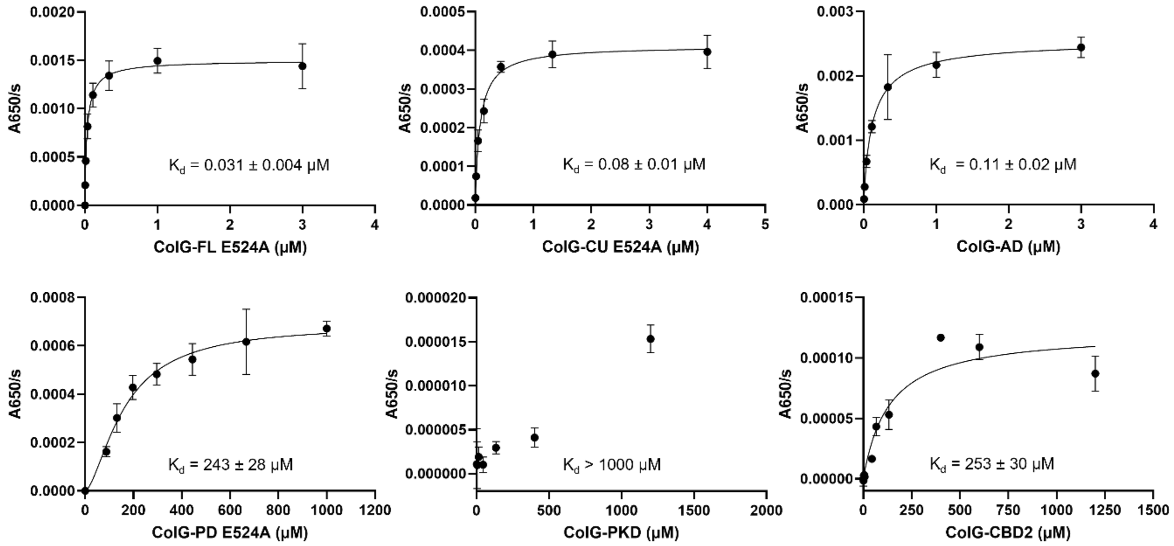


**b** **Type I collagen**

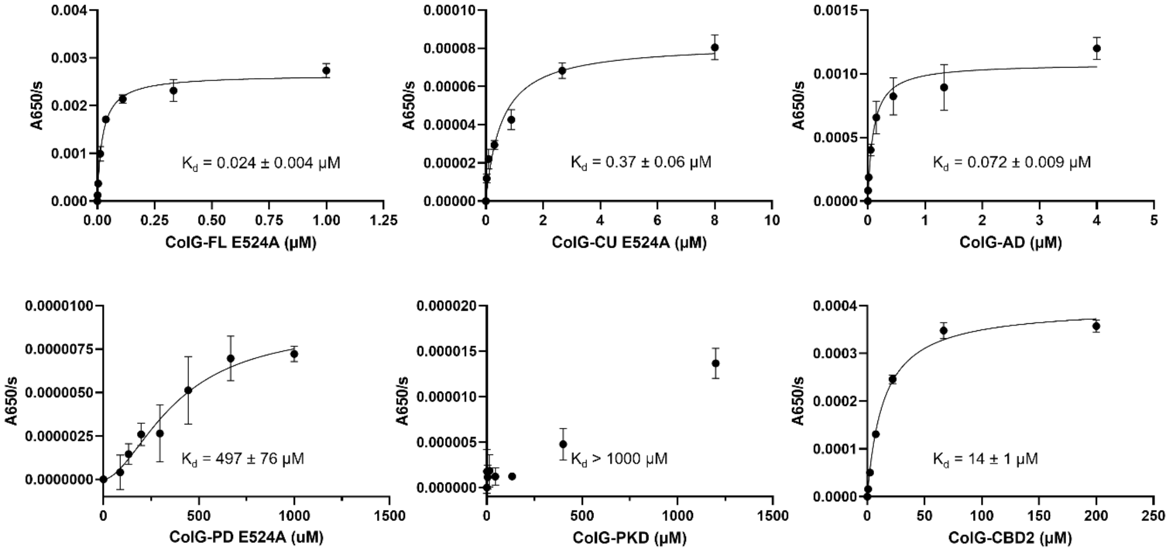


**Fig. S2: ELISA binding assays. a,** Binding curves of ColG variants measured by indirect ELISA on plates coated with denatured type I collagen. Incubation was performed at 37 °C to prevent partial refolding of collagen. **b,** Binding curves of ColG variants to soluble type I collagen. Incubation was performed at 25 °C to prevent collagen unfolding.

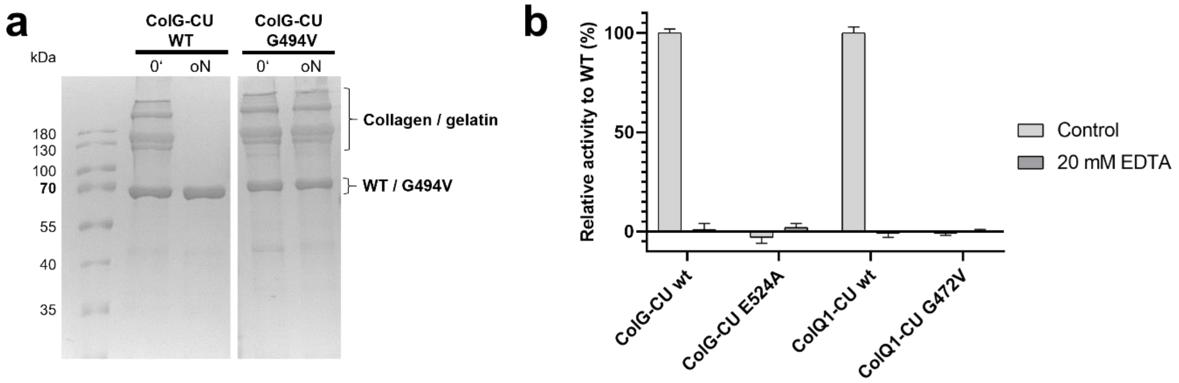


**Fig. S3: ColG-CU G494V, ColG-CU E524A and ColQ1-CU G472V are inactive mutants. a**, 1 mg/ml type I tropocollagen was digested at 25 °C by 4.54 µM ColG-CU variants overnight. Samples were taken at indicated time points and the reaction was stopped by addition of 38 mM EDTA. **b**, ColG-CU and ColQ1-CU variants were incubated with 2 µM FS1-1, a fluorescent-quenched peptide custom-tailored for collagenases, and cleavage was monitored via fluorescence at 328/392 nm at RT.

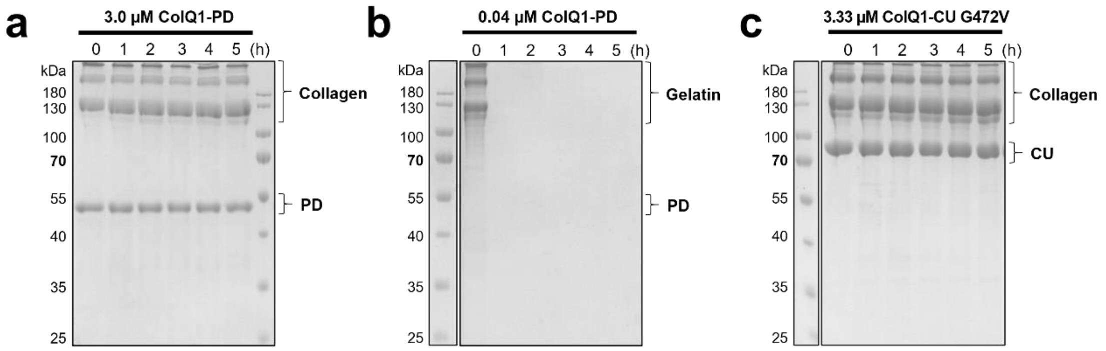


**Fig. S4: ColQ1-PD is an efficient gelatinase, but not collagenase.** ColQ1-PD was incubated with 3.33 µM type I collagen (**a**) or type I gelatin (**b**) at 25 °C or 37 °C, respectively. **c**, ColQ1-CU G472V was incubated with 3.33 µM type I collagen at 25 °C. Samples were taken at indicated time points and the reaction was stopped by addition of 38 mM EDTA.

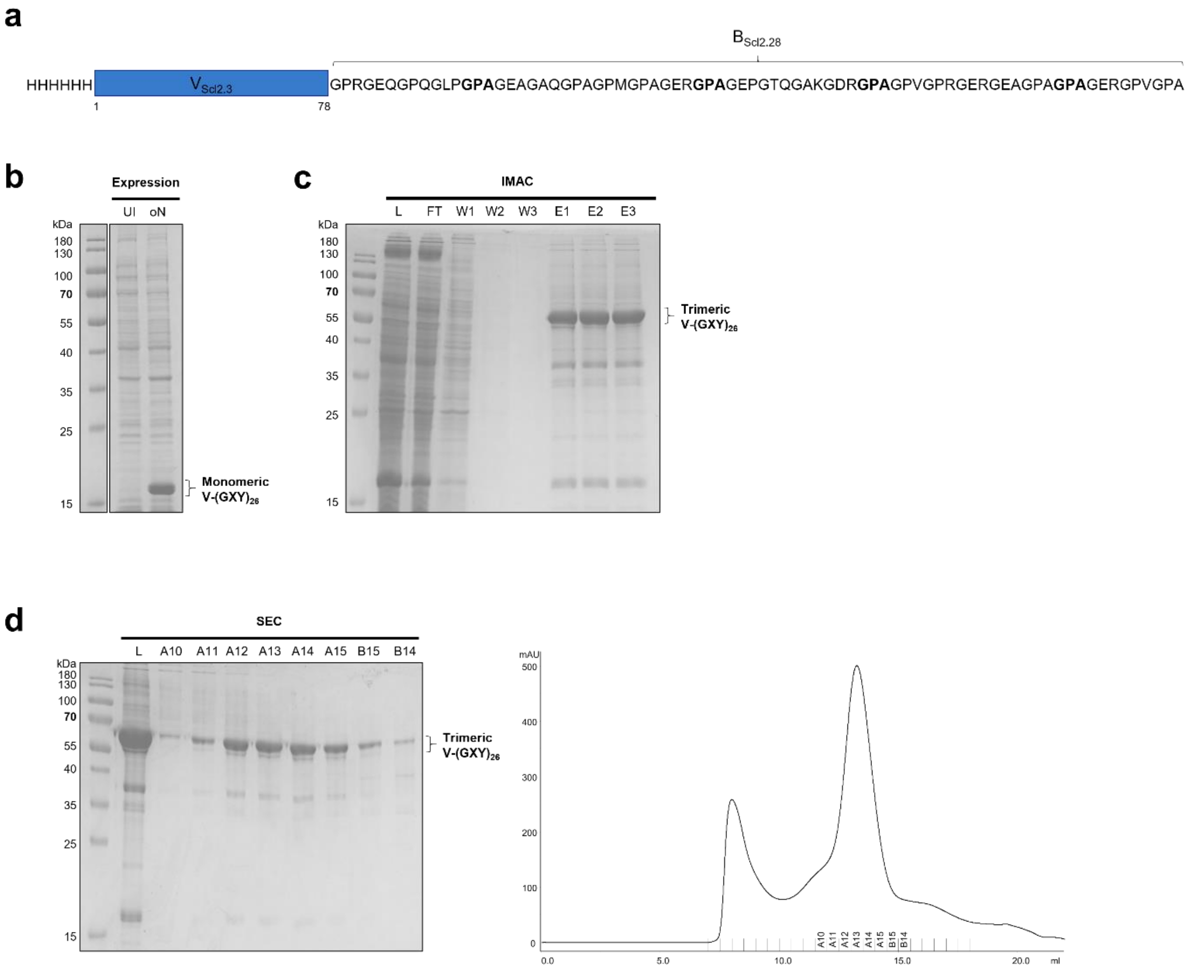


**Fig. S5: Organization and production of V-(GXY)$_{26}$. a**, Scheme of V-(GXY)$_{26}$. Mutations in segment B are indicated in bold. **b**, SDS-PAGE analysis of overnight expression culture at 25 °C in E. coli Nico21 (DE3). Uninduced cell lysate (UI) and cell lysate after overnight expression (oN). **c**, Purification of His-tagged V-(GXY)$_{26}$ via immobilized metal-affinity chromatography (IMAC) using Nickel-Sepharose. **d**, Size-exclusion chromatography (SEC) of His-tagged V-(GXY)$_{26}$ using a Superdex 200 10/300 GL column monitored by SDS-PAGE.

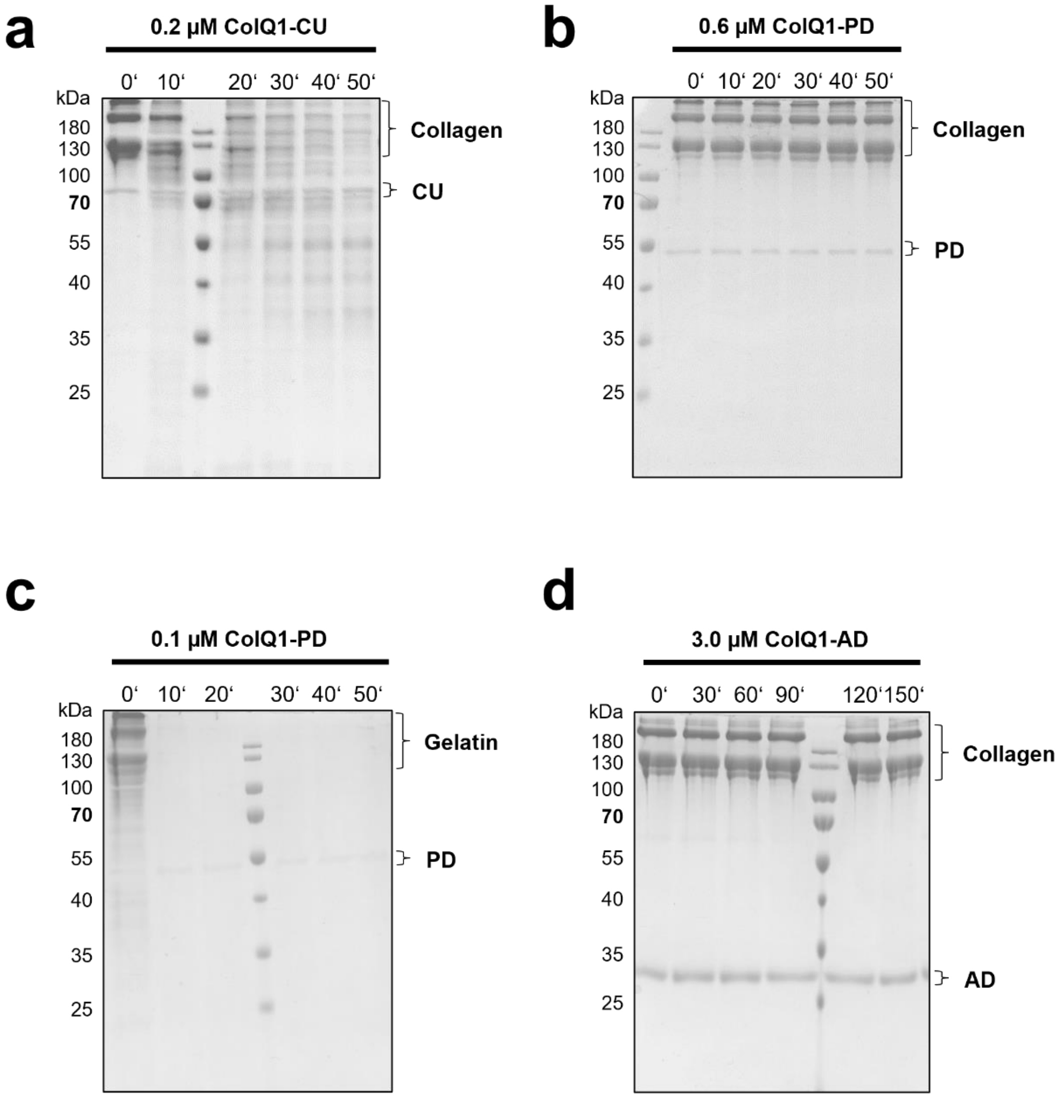


**Fig. S6: ColQ1-CU is an efficient collagenase, whereas ColQ1-PD is an efficient gelatinase, but not collagenase. a,** Degradation of 3.33 µM type I collagen by 0.2 µM ColQ1-CU at 25 °C within 50 min. **b**, Incubation of 3.33 µM type I collagen with 0.6 µM ColQ1-PD at 25 °C did not result in any detectable collagen degradation within 150 min, while in **c** 3.33 µM type I gelatin (generated by heat denaturation of collagen at 95 °C for 5 min) were completely turned over within the first 30 min by 0.1 µM ColQ1-PD at 37 °C. **e**, 3.33 µM type I collagen were stable in the presence of 3.0 µM ColQ1-AD at 25 °C. Reactions were terminated by addition of 35.6 mM EDTA and subjected to SDS–PAGE under non-reducing conditions.

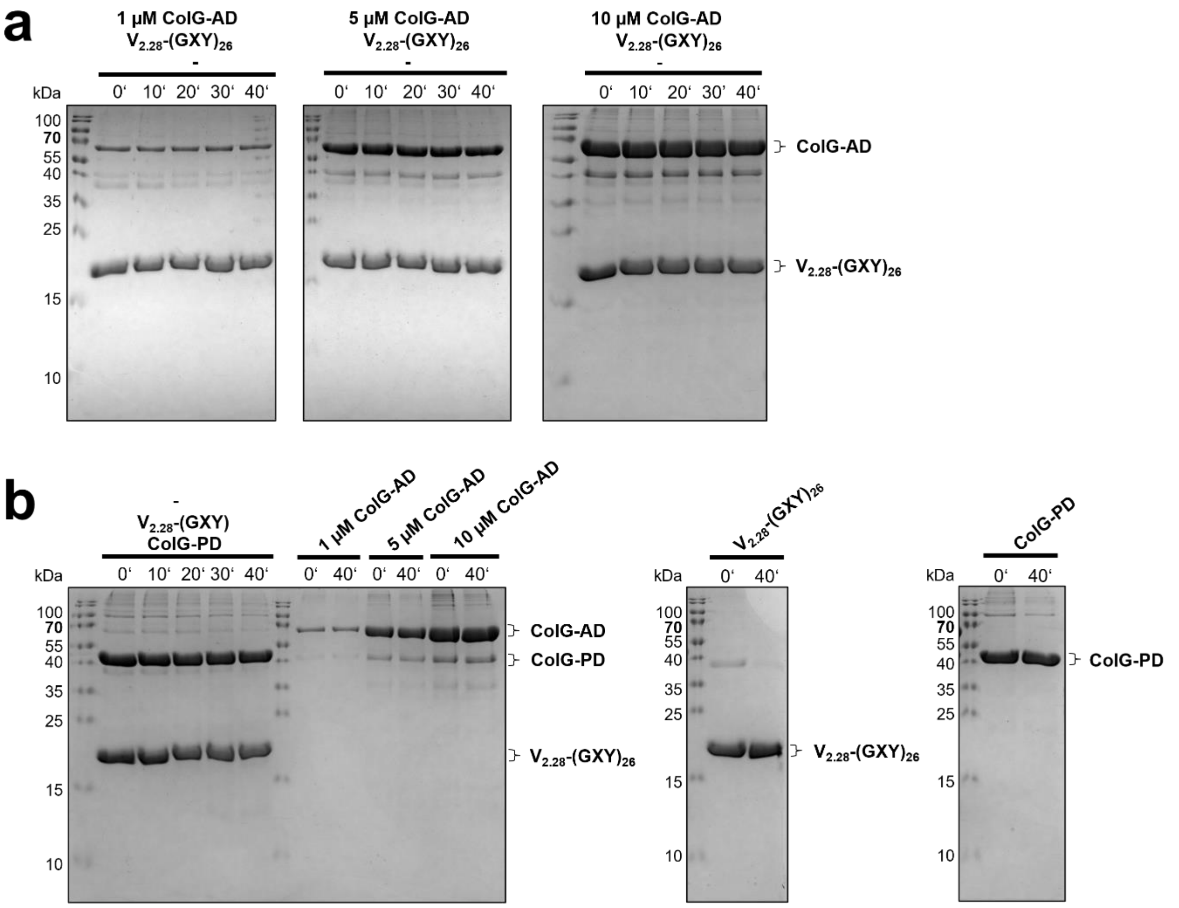


**Fig. S7: The individual AD and PD cannot degrade triple-helical V-(GXY)$_{26}$. a,** Co-incubation of 10 µM V$_{2.28}$-(GXY)$_{26}$ with 1, 5 or 10 µM ColG-AD-MBP. **b**, Co-incubation of 10 µM V$_{2.28}$-(GXY)$_{26}$ with 10 µM ColG-PD and single protein control samples. All reactions were performed at 25 °C. The reactions were stopped by the addition of SDS-loading buffer and analyzed by SDS-PAGE.

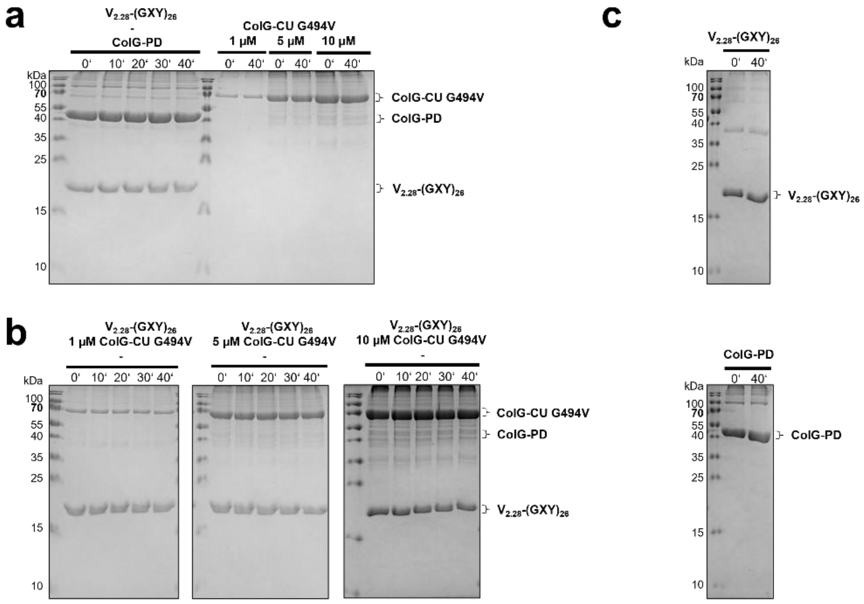


**Fig. S8: Control reactions. a,** Co-incubation of 10 µM $V_{2.28}$-$(GXY)_{26}$ with 10 µM ColG-PD. **b,** Co-incubation of 10 µM $V_{2.28}$-$(GXY)_{26}$ with 1, 5 or 10 µM ColG-CU G494V. **c,** Single protein control samples: 10 µM $V_{2.28}$-$(GXY)_{26}$ and 10 µM ColG-PD. All reactions were performed at 25 °C. The reactions were stopped by the addition of SDS-loading buffer and analyzed by SDS-PAGE.

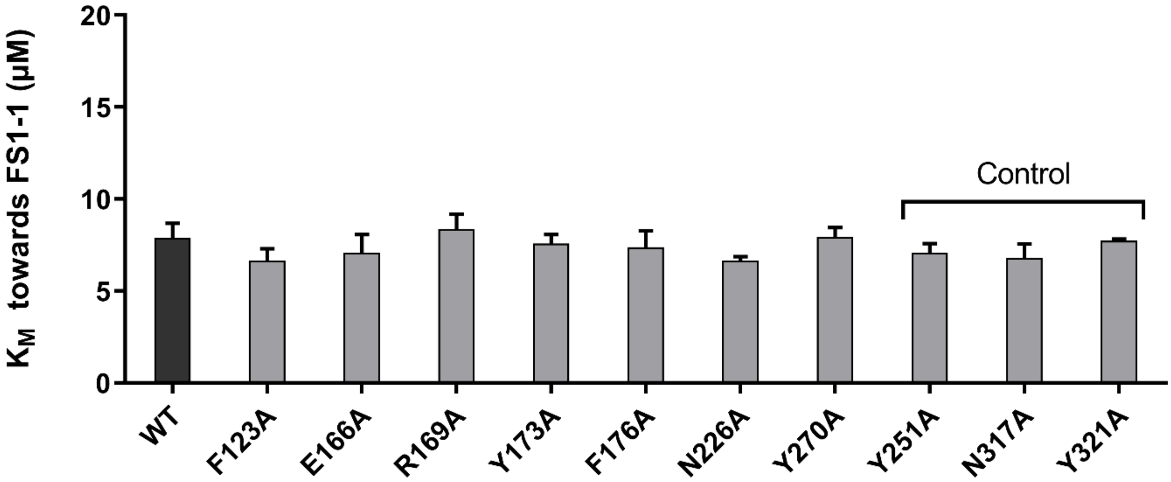


**Fig. S9: Michaelis-Menten constants of ColQ1-CU WT and its single-point mutants towards peptide FS1-1.** Steady state measurements were performed using ColQ1-CU WT and its single-point mutants at 1 nM concentration. The substrate concentration was varied from 0 – 175 µM. Initial velocities were determined using linear regression and the $K_M$ was calculated by non-linear regression using GraphPad Prism 9.1.2 (Graph Pad Software, San Diego, CA, USA). Mutants Y251A, N317A and Y321A were used as controls.

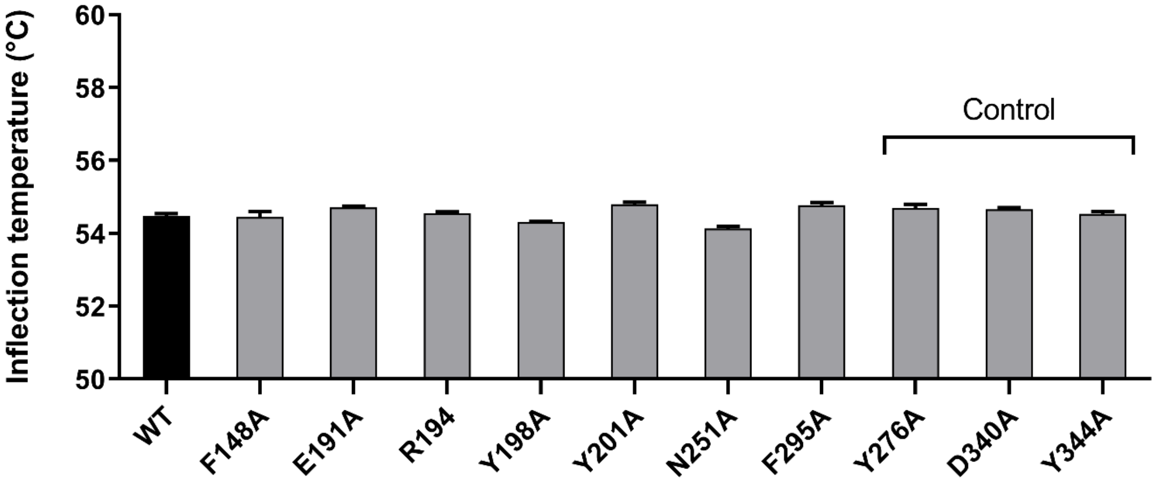


**Fig. S10: Thermal stability of ColG-CU G494 WT and its variants**. Thermal denaturation profiles were measured at 0.1 mg/ml in 15 mM Tris-SO4 pH 7.5, 100 mM NaF and 1 mM $CaCl_2$ using a Tycho NT.6 (Nanotemper, Germany) exploiting the intrinsic fluorescence of tryptophan and tyrosine residues detected at 350 nm and 330 nm. Mutants Y276A, D340A and Y344A were used as controls.

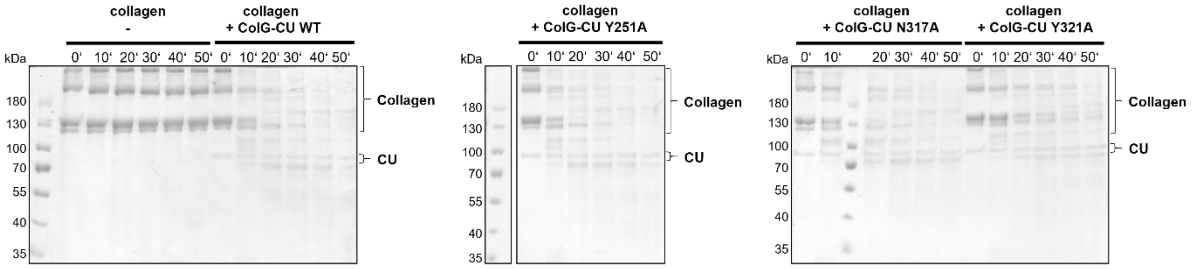


**Fig. S11: Activity of ColQ1-CU WT control mutants towards type I tropocollagen.** 3.33 µM type I collagen were incubated with 0.2 µM ColQ1-CU variants at 25 °C. The reactions were stopped by the addition of 38 mM EDTA and SDS-loading buffer.

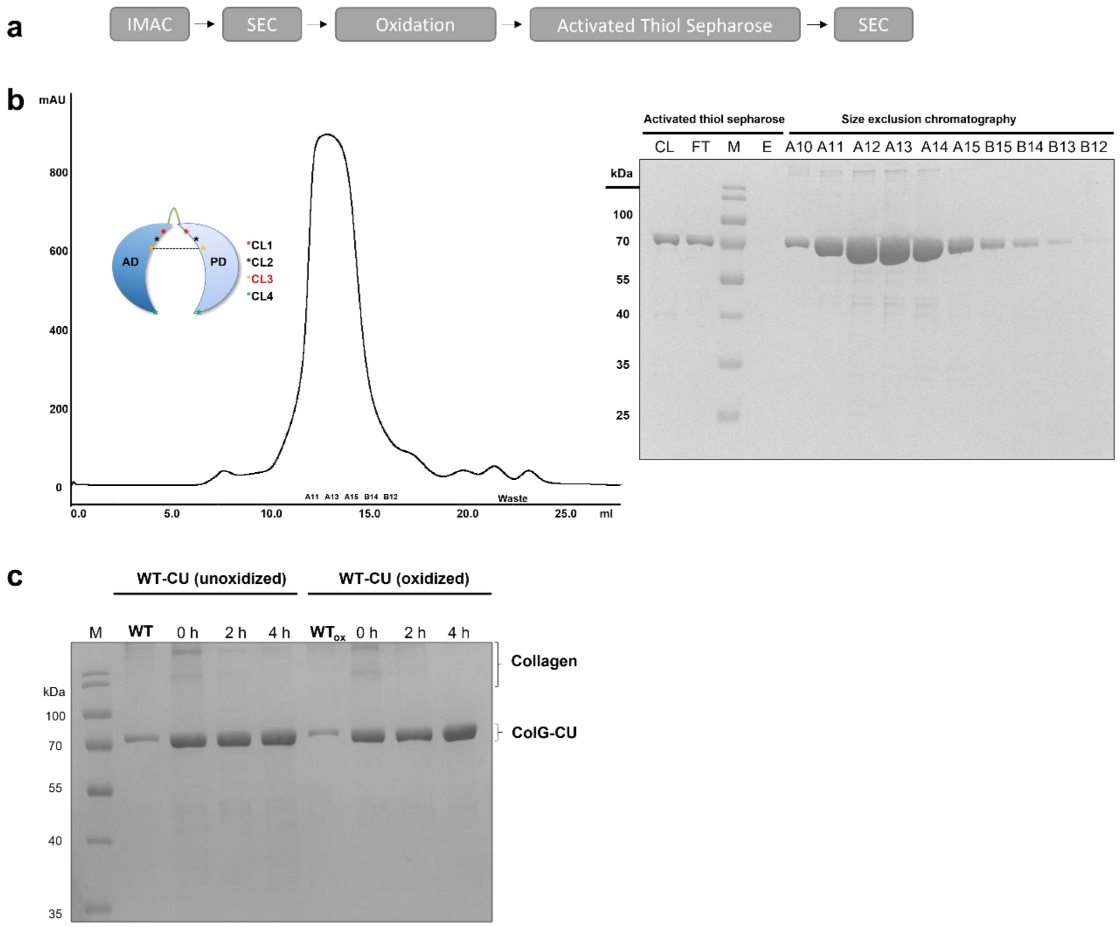


**Fig. S12: Purification of crosslinked ColG-CU mutants. a**, Scheme of production workflow. After metal-affinity purification (IMAC) and a first size-exclusion chromatography (SEC) under reducing conditions, the purified proteins were air-oxidized over 10 d at 4 °C. Non-oxidized species were removed via the Activated Thiol Sepharose resin and misoxidized aggregates were removed by SEC. **b**, Representative example of the final polishing SEC plus corresponding SDS-PAGE analysis including the results of the Activated Thiol Sepharose chromatography. **c**, Oxidation procedure does not affect collagen degradation by ColG-CU WT. ColG-CU WT was stored for 10 d at 4 °C under reducing conditions or air-oxidized over 10 days at 4 °C and then purified by SEC. The purified proteins were tested vs. 1 µM soluble type I collagen at 25 °C.

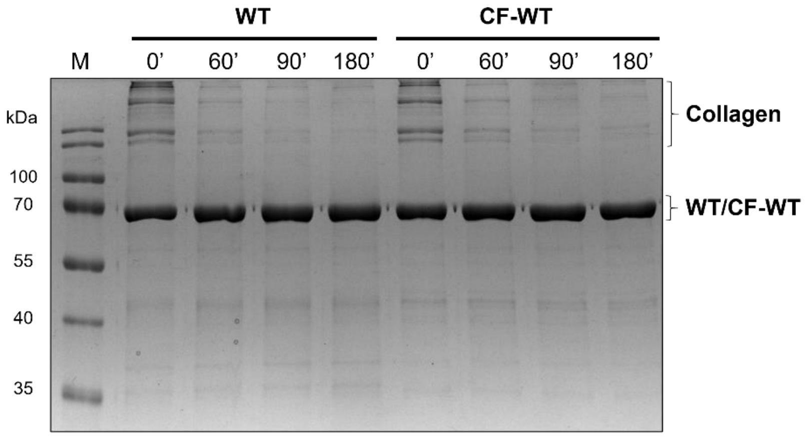


**Fig. S13: CF-WT shows similar collagenolytic activity like ColG-CU WT.** 1 mg/ml type I tropocollagen was digested at 25 °C by 4.54 µM collagenase for up to 4 h. Samples were taken at indicated time points and the reaction was stopped by addition of 38 mM EDTA. The integrity of the collagen fold was verified by co-incubation with 0.83 µM α-chymotrypsin (data not shown).

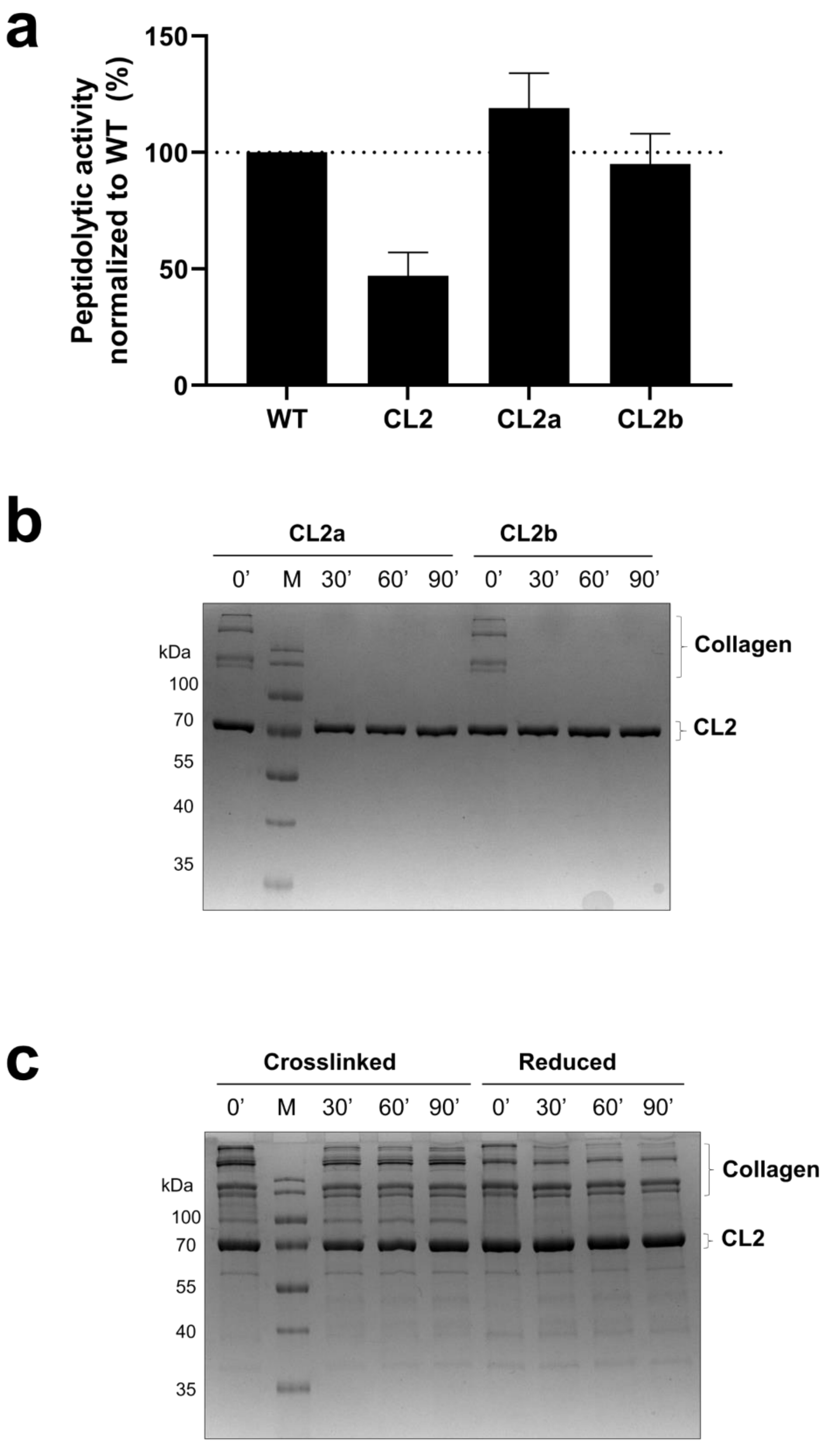


**Fig. S14: Peptidolytic and collagenolytic activity of CL2 and its variants.** **a**, The activities of 16 nM ColG-CU WT, CL2, CL2a and CL2b towards 2 µM quenched-fluorescence substrate. The specific activity of ColG-CU WT was taken as 100%. **b**, Activity of the single-point mutants CL2a and CL2b towards type I tropocollagen over 90 min at 25 °C under non-reducing conditions. **c**, Activity of CL2 towards type I tropocollagen over 90 min at 25 °C under non-reducing and reducing conditions, separated on a non-reducing 12% SDS-PAGE.

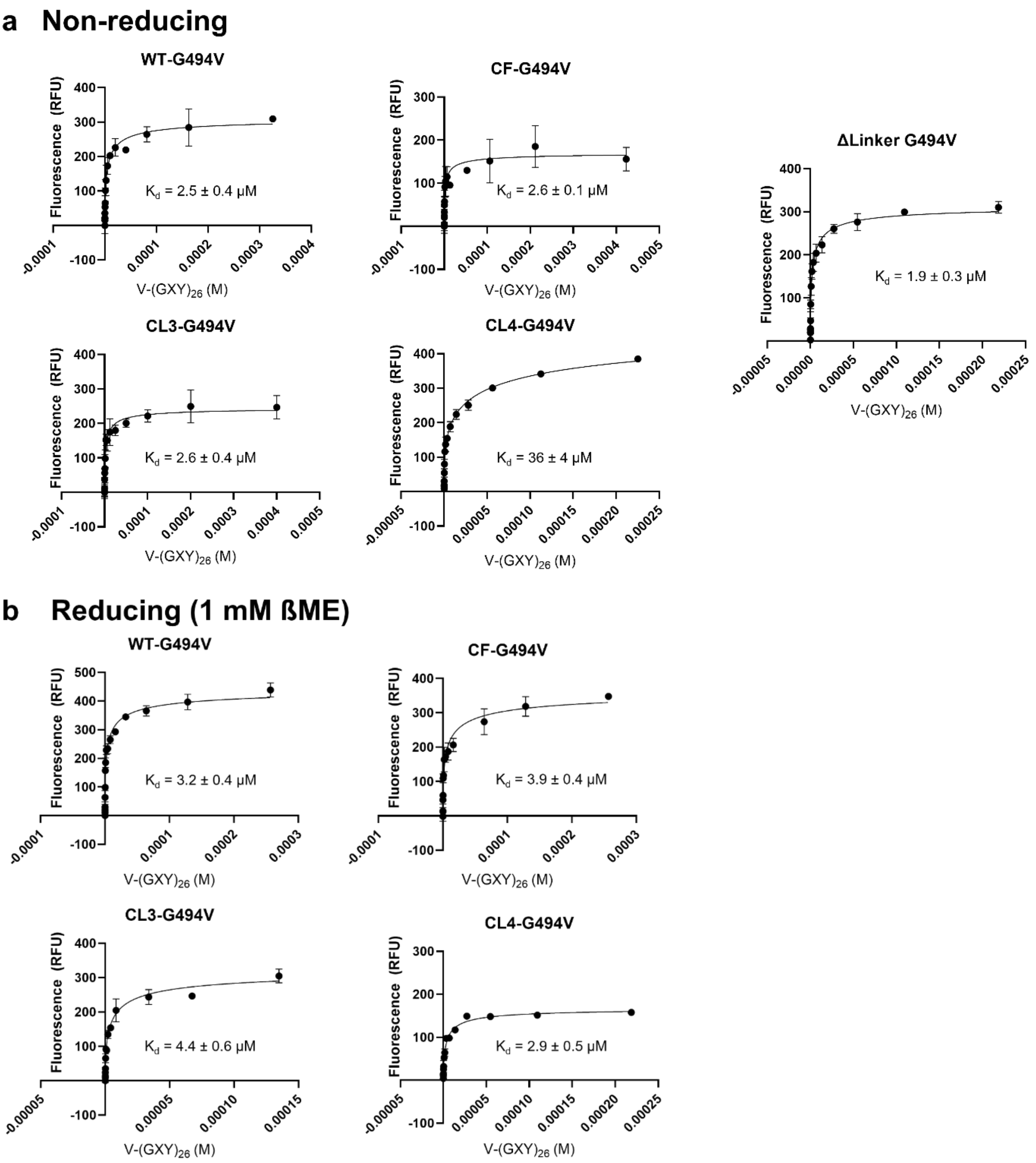


**Fig. S15: MST binding curves.** Binding curves of fluorescently labelled ColG variants towards triple-helical V-$(GXY)_{26}$ measured by microscale thermophoresis using the initial fluorescence signal. Experiments were performed at 22 °C in the absence (**a**) or presence (**b**) of 1 mM ß-mercaptoethanol.

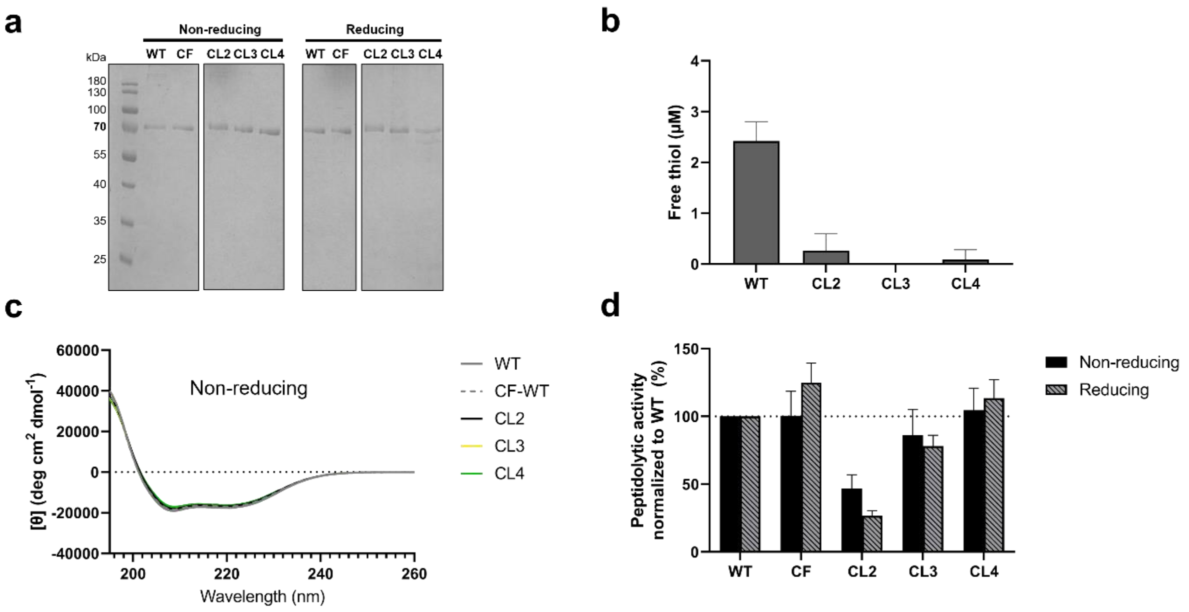


**Fig. S16: Quality control of crosslinked variants CL2-CL4 of ColG-CU.** **a**, SDS-PAGE analysis of ColG-CU WT, CF and the mutants CL2-CL4 under non-reducing and reducing conditions on a 12% polyacrylamide gel. **b**, The presence of free thiols was detected using the thiol-specific fluorochrome 7-diethylamino-3-(4-maleinimidophenyl)-4-methyl coumarin after thermal denaturation of the variants at 60 °C. **c**, CD spectra of ColG-CU WT and the mutants CL2-CL4 in the absence of reducing agent. The data shown are representative of triplicate experiments. **d**, Peptidolytic activity of ColG-CU WT compared to CF and the crosslinked mutants under reducing and non-reducing conditions. 16 nM ColG-CU variants were co-incubated with 2 µM quenched-fluorescent peptide FS1-1 in reaction buffer containing ± 0.5 mM TCEP.

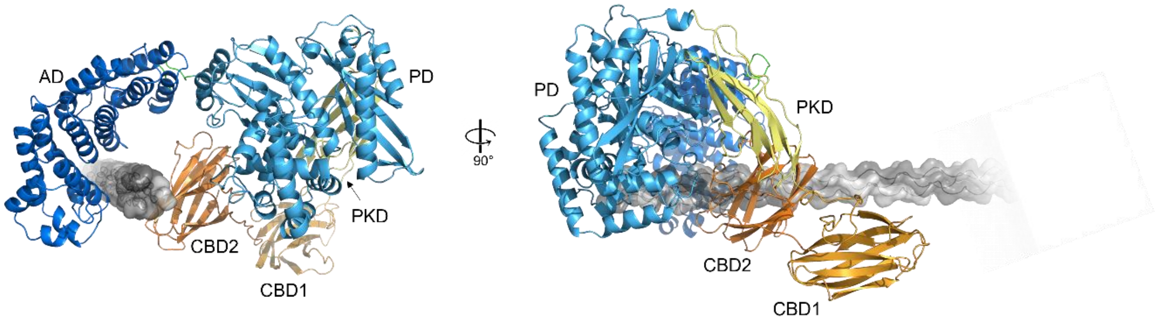


**Fig. S17: Docking model of ColG-FL on collagen triple helix.** Ribbon and surface representation of ColG-FL and a mini collagen triple helix composed (GPP)60. The structure models were generated with Alphafold2 and then docked with HADDOCK 2.4, an information-driven docking tool. Although binding residues for AD, CBD1 and CBD2 were defined, the docking only positioned the AD and CBD2 on the triple helix. The latter docking is in line with the findings that i) the linker between CBD1 and CBD2 does not allow for both domains to bind simultaneously to the same tropocollagen molecule[3] and ii) that the CBD2 domain exhibits the higher affinity to collagen than CBD1 [4]. The structures of the individual CU, PKD and CBD2 of the docked ColG-FL structurally superimpose well with their respective crystal structures (RMSD = 1.405 / 0.747 / 0.588, PDB: 2y50 / 2y72 / 4HPK calculated using Pymol[5]). The same color code for ColG as in Fig. 1 is used.

**Table S1: Dissociation constants ($K_d$) of ColQ1-CU G494V mutants towards type I gelatin and type I collagen determined by ELISA.**

| | **Gelatin** $K_d$ (mean ± SD) (µM) | **Collagen** $K_d$ (mean ± SD) (µM) |
|---|---|---|
| **WT** | 1.6 ± 0.4 | 1.6 ± 0.2 |
| **F123A** | 1.6 ± 0.3 | nbd |
| **E166A** | 5.0 ± 0.6 | nbd |
| **R169A** | 2.8 ± 0.4 | 0.17 ± 0.04 |
| **Y173A** | 24 ± 6 | nbd |
| **F176A** | 1.2 ± 0.2 | 0.08 ± 0.01 |
| **N226A** | > 150 | > 150 |
| **Y270A** | 4.2 ± 0.5 | 0.19 ± 0.06 |
| **Y251A** | 2.5 ± 0.2 | 2.8 ± 0.6 |
| **N317A** | 2.3 ± 0.1 | 2.8 ± 0.5 |
| **Y321A** | 2.5 ± 0.1 | 4.0 ± 0.4 |

The apparent dissociation constant $K_d$ is given as mean value of three independent experiments ± standard deviation. nbd, no binding detected.

### 2. Experimental section

**Rational design and production of crosslinked ColG-CU variants.**

To ensure efficient disulfide-bridge formation, we were looking for non-conserved residues on the inner-facing surfaces of the AD and PD. We generated a model of (semi)-closed conformations of ColG-CU based on PDB entry 2y50 using PYMOL software [5] and identified two residue pairs Y280/Q512 (mutant CL2), and E294/T483 (mutant CL3) for the introduction of cysteines in the upper half of ColG-CU, located at varying distances from the linker region (**Fig. 8a-b**). For mutant CL4, cysteines were introduced in a loop of ColG-PD and just before the N-terminus, in order to crosslink the ColG-CU at the tips of the AD and PD domains, locking the CU in a closed conformation. The mutants CL2-CL4 were generated on the basis of a cysteine-free ColG-CU (C218S/C262S) (mutant CF).

All ColG-CU constructs yielded over 20 mg of homogenous monodisperse protein after purification and oxidation from two liters of *E. coli* cell culture. They migrated with an apparent molecular mass of 79 kDa on a denaturing non-reducing SDS-PAGE gel and were estimated to be approximately 95% pure (**Fig. S16a**). SDS-PAGE analysis revealed that there were negligible amounts of oligomeric forms of the crosslinked mutants, indicating the robustness of the crosslinking approach, and we confirmed that the oxidation process did not negatively affect the collagenolytic activity *per se* (**Fig. S12c**).

The presence of the disulfide linkage was confirmed by a thiol quantitation assay (**Fig. S16b**). All crosslinked ColG-CU variants were tested at 1 µM concentration. ColG-CU WT which harbors two buried cysteines in the AD was used as positive control and we could confirm the presence of the disulfide bonds in CL2 to CL4. Non-reducing CD spectroscopy analysis showed that all mutants had a secondary structure similar to ColG-CU WT, suggesting that the formation of the disulfide bond did not compromise the overall fold (**Fig. S16c**). Finally, we compared the activity of the mutants towards a small quenched-fluorescence peptide substrate to the activity of ColG-CU WT to confirm the proper folding of the PD in the crosslinked variants **Fig. S16d**). The removal of the two native cysteines in the AD in CF did not compromise its peptidolytic activity and collagenolytic activity (**Fig. S13**). In CL3 and CL4, the additional introduction of the cysteines for crosslinking also did not inhibit peptide hydrolysis in the reduced state (86 ± 19%, 105 ± 16%, respectively) and in the oxidized state, when the crosslink was established (78 ± 8%, and 113 ± 14%, respectively). However, mutant CL2 showed a notably reduced substrate turnover in both states (47 ± 10% and 27 ± 4% residual activity under non-reducing and reducing conditions, respectively).

**Model generation for the complex of ColG-FL with a mini collagen triple-helix using protein–protein docking**

A full-length model of ColG was generated using Alphafold2[6]. MMseqs2 and HHsearch with the PDB100 option were used to generate templates, thereby integrating the available structural information of the single ColG domains into the Alphafold predictions. The resulting five relaxed models were manually curated. The Alphafold model which positioned the CBD domains correctly behind the CU was used for further docking. Alphafold2 was also used to generate a mini collagen triple helix based on the sequence $(GPP)_{60}$. Protein-protein docking of ColG-FL to the $(GPP)_{60}$ triple helix was performed using the HADDOCK 2.4 server[7,8], as this docking method allows for the definition of known binding sites as input for the docking procedure. The active site/interface residues of the AD (F148, E191, Y198, N251), CBD1 (Y950, H958, F983, Y985, H987) and CBD2 (L1034, S1038, Y1080, L1102, Y1104, Y1106) were given as input. In addition, the linker regions between the AD and PD (389-397) and between CBD1 and CBD 2 (999-1004) were defined as fully flexible. The resulting complex models were manually curated.
